# Dorsal Attention Network as a Convergent Hub of Diverse Non-pharmacological Interventions Preserving Functional Gradients and Cognition in Aging

**DOI:** 10.1101/2025.07.26.666919

**Authors:** Xinhu Jin, Wei Tang, Zhiwei Zheng, Pengyun Wang, Xinyi Zhu, Xiaoyu Cui, Shufei Yin, Zeping Lv, Haiqun Xie, Rui Li, Juan Li

## Abstract

Non-pharmacological interventions (NPIs) in aging neuroscience have largely focused on intervention-specific regional effects, with limited understanding of generalizable network-level mechanisms. Here, adopting a previously unexplored gradient-based perspective of functional brain organization, we analyzed an NPI dataset involving four interventions in older adults (training/control group: n = 112/59). NPIs led to strengthened intra-network functional integration and maintained macroscale gradient architecture. Virtual lesion analyses identified the dorsal attention network (DAN) as a key contributor to gradient maintenance. Critically, enhanced post-intervention DAN connectivity was associated with maintained gradient structure and improved global cognition. These findings establish a unifying framework in which the DAN acts as a convergent hub through which diverse NPIs preserve functional brain organization and attenuate cognitive aging.

---

Population aging has emerged as a global public health challenge, driving an urgent need to address age-related cognitive decline and neurodegenerative disorders such as Alzheimer’s disease (AD). Non-pharmacological interventions (NPIs), defined as structured, nonchemical therapies grounded in theoretical frameworks^1^, are increasingly recognized for their potential to delay cognitive deterioration and reduce dementia incidence in older adults^2–5^. Unlike pharmacological approaches, NPIs demonstrate favorable safety profiles, higher patient acceptance, and greater scalability across diverse clinical settings^4^. Current evidence highlights promising NPIs for mitigating cognitive decline during healthy and pathological aging, including cognitive intervention (CI), physical intervention (PI), non-invasive brain stimulation (NIBS), and multimodal intervention integrating the approaches above.

NPIs could enhance neuroplasticity in aging populations^6^, defined as the brain’s capacity for structural and functional adaptation to environmental demands^3^. Thus to counteract age-related decline, different types of interventions such as CI^7^, PI^8^, and music training^9,10^ promote neuroplasticity by preserving youth-like patterns of brain activity and functional connectivity. However, three critical constraints impede progress in the current field: First, most studies employ single-intervention designs with small cohorts, restricting the generalizability of findings; Second, existing neuroimaging evidence predominantly focuses on regional morphological or functional activation/connectivity changes, exhibiting substantial inter-study variability while failing to establish a unified explanatory framework; Last, research on neural mechanisms across different types of NPIs remains siloed, with no elucidation of core shared brain mechanisms transcending intervention modalities. Future advances require identifying shared neural mechanisms underlying NPIs at the brain system-level, which are essential to both decipher late-life neuroplasticity and refine theoretical models of how targeted interventions remodel aging brain function.

Age-related network dedifferentiation, the key feature of aging characterized by reduced within-network and increased between-network connectivity^11,12^, provides a rational perspective from functional network organization for deciphering the neural mechanisms of NPIs against aging^2,3^. Functional gradients, a novel metric characterizing the brain’s macroscale hierarchical organization from primary to higher-order functional networks along a principal axis^13^, have established a new paradigm for identifying the core feature of brain architecture in adolescents^14^, adults^13^, the aging population^15^, as well as connectome gradient dysfunctions between atypical and normal groups^16,17^. Although used to characterize functional network changes or differences in observational cohorts, this method remains unexplored for mapping intervention-induced effects in the context of anti-aging research by capturing organizational changes of effective interventions. Moreover, cognitive training induces neuroplastic enhancements that improve both task-specific performance (near-transfer effects) and cross-domain abilities (far-transfer effects) via functional reorganization of the dorsal attention network (DAN)^18^. A systematic umbrella review and meta-meta-analysis confirms the effectiveness of exercise for improving general cognition, memory, and executive function^19^, though the neuroprotective mechanisms underpinning these far-transfer effects in healthy brain aging remain poorly defined. Multimodal interventions promote the recruitment of compensatory neural resources, further supporting core network engagement to meet task demands^20^. Hence, beyond intervention-specific benefits, a convergent functional network may exist via which each type of intervention effectively attenuates age-related cross-domain cognitive decline and preserves younger brain organization.

In the present study, we attempt to expand the theoretical framework of non-pharmacological intervention and neuroplasticity in older adults against aging by mapping pre-to-post-intervention changes in functional architecture and then further revealing intervention-general network transcending intervention-specific neural changes across diverse NPIs. Adopting four different types of intervention from our lab (Cognitive Training; Square Dance; Combined Intervention of Aerobic Exercise/Video Game; Multimodal Intervention of Cognitive Training/Tai Chi/Group Counseling), we constructed an NPI dataset by integrating older adults in the control group (CG) and training group (TG) both pre- and post-intervention, and first identified pronounced behavior (global cognition) and brain (functional connectivity strength and functional gradient based on resting-state functional magnetic resonance imaging) changes after intervention between CG and TG. According to the “last in, first out” model^21,22^, association areas not only show mirroring of healthy developmental and aging processes but also demonstrate heightened sensibility to etiologically distinct clinical disorders linked to abnormal aging trajectories (AD) and effective NPIs counteracting aging-related deterioration^2,3,22,23^. Therefore, we hypothesized higher-order networks such as DAN, default-mode network (DMN), or frontoparietal network (FPN) as potential shared networks across heterogeneous NPIs and tested them via follow-up virtual lesion analyses^24^, which excluded network regions in post-intervention TG and then rederived functional gradients. Highlighting the importance of convergent networks in cognitive functioning, global cognition enhancement was only linked with increased functional connectivity strength within DAN in TG after intervention, along with the gradient architecture maintenance. In summary, dorsal attention network is the core convergent hub underlying diverse non-pharmacological interventions preserving gradient organization and global cognition in aging. These findings will provide important theoretical and empirical evidence for promoting healthy brain aging and the prevention as well as intervention of neurodegenerative disorders.

## Results

### The primary outcome demonstrates a significant intervention effect

This study constitutes a pooled analysis of previously registered NPIs (see Methods; Supplementary Table S1). The final analysis included 112 TG participants and 59 CG participants (Fig. 1). Detailed demographic characteristics and neuropsychological test performances are presented in Supplementary Table S2.

**Fig. 1.**
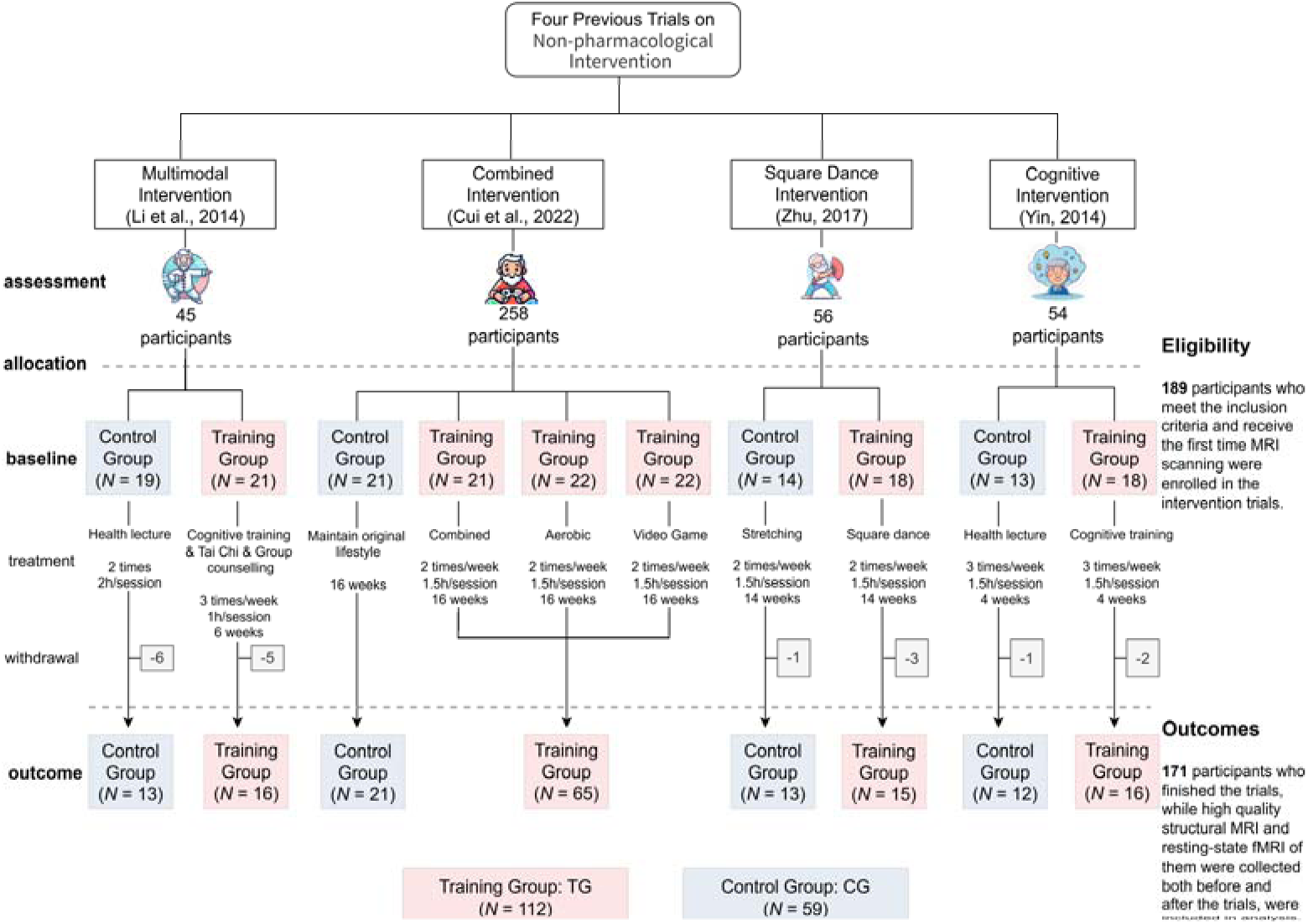
Intervention flowchart. This flowchart illustrates the recruitment and exclusion process of four registered non-pharmacological interventions (NPIs) we used. Participants were excluded if they lacked a final MRI scan or had low imaging quality. The training group (TG) included participants from the multimodal, combined, aerobic, video game, square dance, and cognitive intervention groups (labeled in red), while the control group (CG) comprised participants from the health lecture, original lifestyle maintenance, and stretching exercise groups (labeled in blue).

As shown in Fig. 2a, a significant interaction effect of group (TG vs. CG) × period (pre- vs. post-intervention) was found for all participants measured with the Montreal Cognitive Assessment (MoCA) as the primary outcome (TG/CG: *n* = 97/46; *F*(1, 141) = 4.31, *p* = 0.040, partial η^2^ = 0.030). Simple main effects analysis further revealed that the interaction effect was driven by increased MoCA scores in TG after intervention (post-pre: 0.95, 95% CI = 0.59-1.31, *p* < 0.001, partial η^2^ = 0.162), with no significant change observed in CG (post-pre: 0.28, 95% CI = −0.24-0.81, *p* = 0.286, partial η^2^ = 0.008). Consequently, heterogeneous NPIs significantly enhanced TG participants’ global cognition, suggesting the existence of far-transfer effects.

**Fig. 2.**
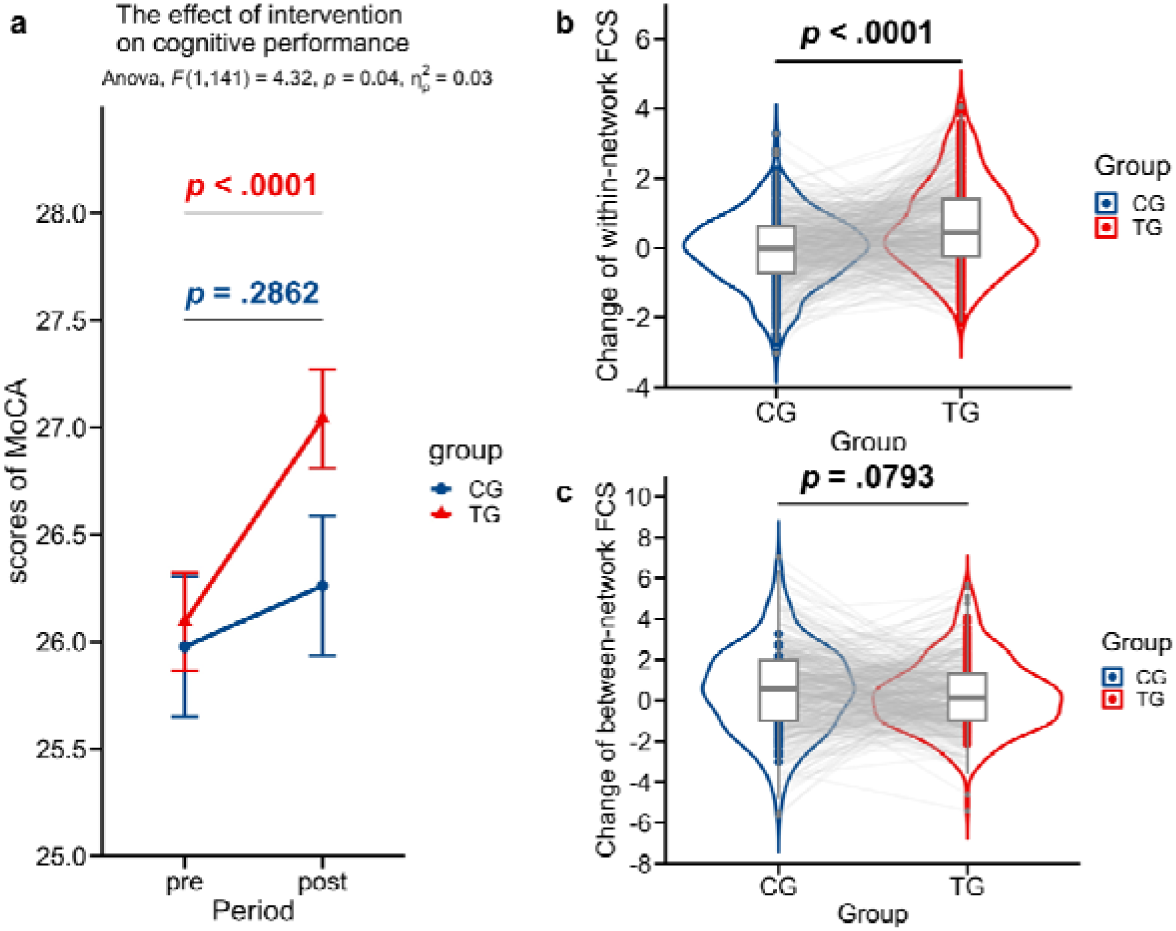
Intervention effects on primary outcome and brain connectivity. Significant intervention effects were observed on (a) MoCA and (b) within-FCS. Conversely, no significant group difference was found in (c) between-FCS. These findings revealed clear intervention effects on global cognition and intra-network integration, providing potential neural evidence for the enhanced primary outcome in TG. Higher values indicate better global cognition or increased FCS after intervention trials. Error bars represent the standard error of the means. Gray lines link values of identical brain parcels across two groups. MoCA, Montreal Cognitive Assessment; TG/CG, training/control group; FCS, functional connectivity strength.

We also calculated the interaction effects for secondary outcomes, including measures of specific cognitive functions: digital span-forward/backward (DSF/DSB), Paired Associative Learning Test (PALT), and verbal fluency (Supplementary Fig. S1). For all participants who have completed DSF/DSB tests, no significant interaction effect was observed (Both *p* ≥ 0.770, partial η^2^ < 0.001). In parallel, the interaction effect for PALT was significant (*p* < 0.001, partial η^2^ = 0.100), while TG participants exhibited a trend toward improved verbal fluency performance (*p* = 0.006) though no interaction effect was found (*p* = 0.310, partial η^2^ = 0.006), consistently showing multidomain intervention effects despite methodological heterogeneity.

### Functional connectivity strength within the same network serves as a target of diverse NPIs

To investigate neural mechanisms underlying NPIs, we constructed parcel-parcel resting-state functional connectivity (rsFC) matrices for all participants (*n* = 171) based on the Schaefer parcellations with 400 parcels^25^. All 400 cortical parcels were mapped into 7 networks^26^. We separately generated group-level Fisher-z transformed FC (FCz) matrices that harmonized the multi-site effect for TG (*n* = 112) as well as CG (*n* = 59) and then computed both within-network functional connectivity strength (within-FCS; sum of FCz between the parcels belonging to the same network) and between-network functional connectivity strength (between-FCS; sum of FCz between the parcels belonging to different networks) for each parcel, where FCS reflects the global associations of a given parcel with the rest brain parcels (Methods).

Comparative analysis of pre-to-post intervention FCS changes between the TG and CG revealed an intervention-induced enhancement of within-FCS in TG (TG-CG: 0.60, 95% CI = 0.43-0.78, *p* < 0.001, Cohen’s *d* = 0.403; Fig. 2b), whereas between-FCS changes showed no significant group difference but a decrease trend in TG (TG-CG: −0.25, 95% CI = −0.53-0.03, *p* = 0.079, Cohen’s *d* = −0.088; Fig. 2c).

These findings demonstrated that NPIs might primarily modulate cognitive function by strengthening intra-network functional integration, suggesting a mechanism rooted in optimized local information processing within specialized functional circuits.

### Diverse NPIs counteract age-related reorganization by maintaining brain hierarchical gradient organization

Age-related network dedifferentiation may contribute to disruptions in macroscale hierarchical organization, as reflected by gradient architecture^15,27^. Thus, we hypothesized that NPIs may exert intervention effects by enhancing within-network FCS in core target networks, thereby helping maintain hierarchical organization. To verify this hypothesis, we retained the top 10% functional connections per parcel in group-level FCz matrices to enforce sparsity. Functional connectivity gradients were then computed using diffusion map embedding^28^, allowing the decomposition of individual-level connectivity matrices into a lower-dimensional space. The resulting gradients reflect spatially dissociable patterns of cortical connectivity, ordered by the variance explained in the initial FCz matrix (Methods).

Before intervention, both TG and CG exhibited highly similar gradient patterns (TG1 and CG1), characterized by primary gradient (G1) that extended from unimodal (somato/motor and visual) to transmodal (association) areas (S-A axis), and secondary gradient (G2) spanning the somato/motor-visual axis. However, after the interventions, TG participants (TG2) retained this gradient organization, whereas CG participants (CG2) exhibited a swapped gradient topology. Specifically, primary gradients in TG1, CG1, and TG2 closely matched the secondary gradient in CG2 (Fig. 3a-d). Moreover, we used the gradient pattern of TG1 as a reference template and mapped the spatial distribution of gradients across the 400 parcels for each group at both pre- and post-intervention periods (Fig. 3e). All results revealed a consistent gradient organization in TG1, CG1, and TG2, indicating the maintenance of intrinsic cortical hierarchy following NPIs. In contrast, only CG2 (lacking effective intervention) exhibited a gradient pattern deviation from the original hierarchical organization.

**Fig. 3.**
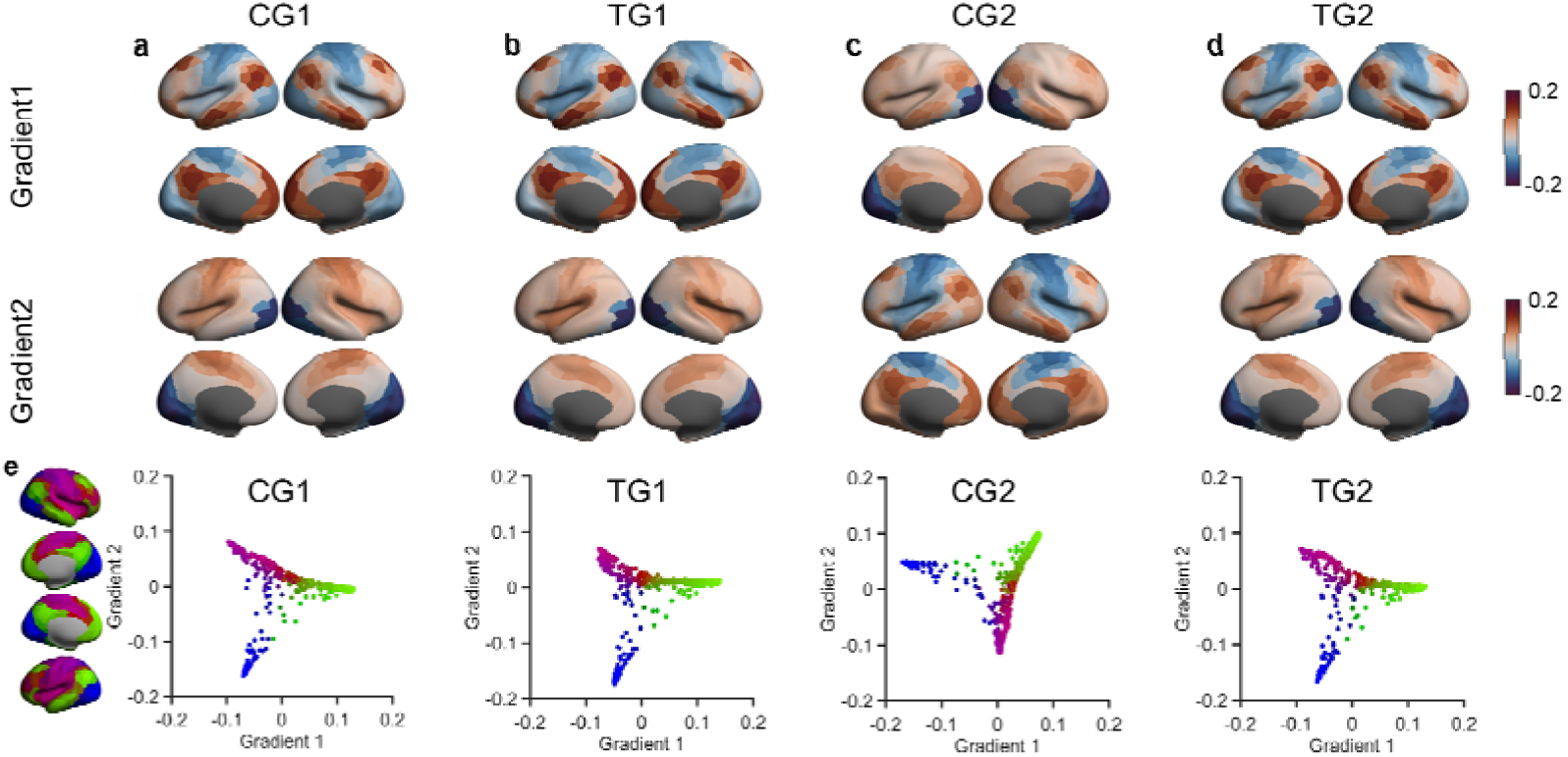
Non-pharmacological interventions maintain functional connectivity gradients. The primary and secondary gradients of functional connectivity in (a) CG1, (b) TG1, (c) CG2, and (d) TG2. Before intervention, both TG1 and CG1 showed a consistent primary gradient (G1) from unimodal (somato/motor and visual) to transmodal (association) areas, and a secondary gradient (G2) separating somato/motor and visual areas. After intervention, TG2 retained the gradient organization, while CG2 (lacking effective NPIs) exhibited a swapped gradient pattern. Gradient colours (blue-red) represent gradient values. (e) Using TG1 as a reference, gradient values of 400 cortical parcels were exhibited with scatter plots, showing consistent gradient patterns in TG1, CG1, and TG2, but a clear reversal pattern in CG2. TG1/CG1, pre-intervention training/control group; TG2/CG2, post-intervention training/control group; NPIs, non-pharmacological interventions.

### A core role for the dorsal attention network in sustaining hierarchical brain organization after interventions

Having established that NPIs maintained hierarchical cortical organization against aging in the TG, we next sought to identify the core functional network(s) mediating this neuroprotective effect. Here, we implemented a new analysis procedure called “virtual lesion”^24^ by directly excluding network parcels from the group-level TG2 FCz matrix (*n* = 112) and rederived the functional gradients to investigate which network defined by Yeo et al.^26^ played a central role in sustaining the gradient architecture against brain aging. The dorsal attention network (DAN), known for its role in top-down attentional control^18^, emerged as a crucial determinant across diverse NPIs in maintaining the cortical gradient organization. Specifically, when DAN was selectively removed from TG2 (Fig. 4b), the rederived primary gradient now closely resembled the primary gradient of CG2 (Fig. 4a; *r* = 0.96, *p*_spin_ < 0.001, 1000 permutations). Similarly, the rederived secondary gradient in DAN-dropped TG2 shifted to match CG2’s secondary gradient (*r* = 0.96, *p*_spin_ < 0.001, 1000 permutations). Scatter plots confirmed and demonstrated that DAN-dropped TG2 exhibited a gradient architecture similar to CG2 (Fig. 4c-d), supporting our hypothesis that the general intervention-driven optimization of DAN plays a core role in maintaining hierarchical cortical organization, thus mitigating brain aging.

**Fig. 4.**
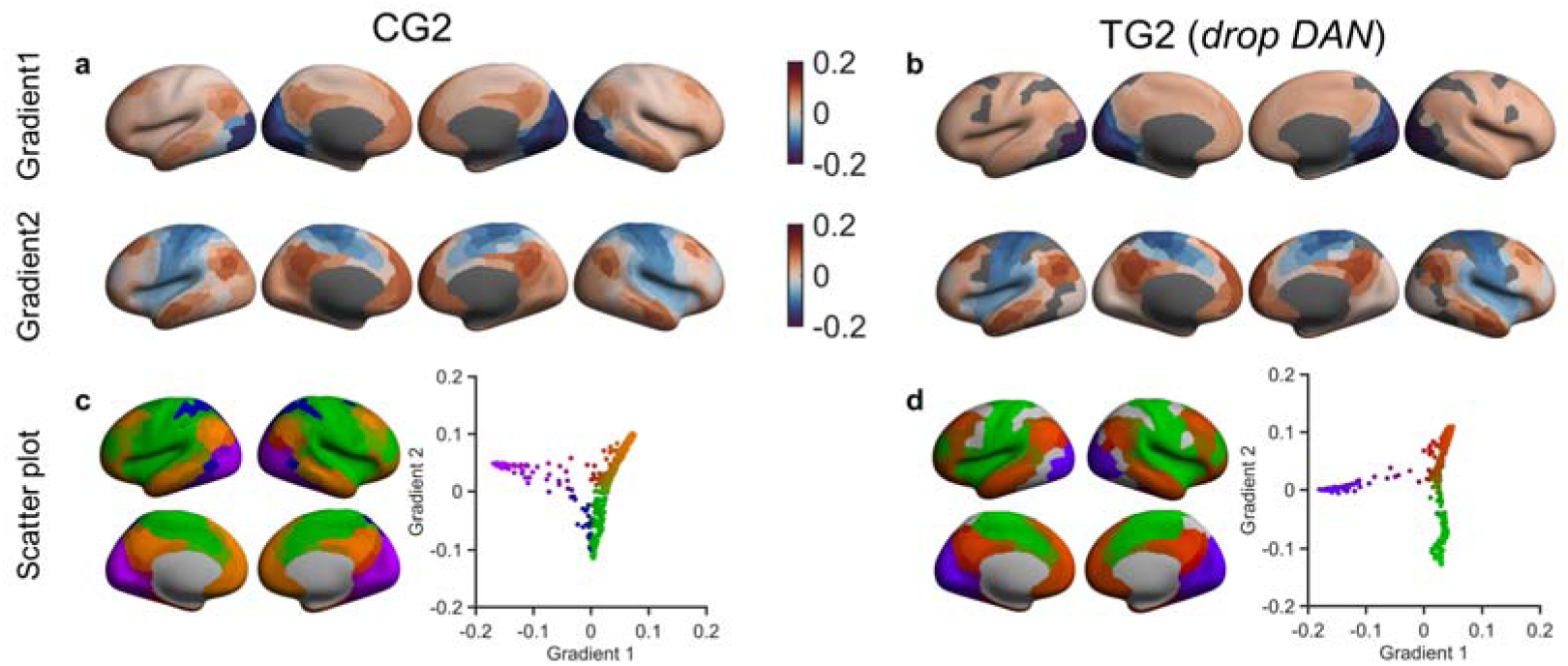
The dorsal attention network plays a core role in maintaining gradient architecture against brain aging. Primary and secondary gradients of (a) CG2 and (b) DAN-dropped TG2. Results revealed that removing DAN from TG2 disrupted its original gradient organization, shifting it to resemble the CG2 pattern, which reflected the normal aging process without intervention. The primary and secondary gradients of DAN-dropped TG2 closely matched those of CG2, whereas no such similarity was found between DAN-dropped TG2’s primary gradient and CG2’s secondary gradient, or vice versa. Gradient colours (blue-red) represent gradient values. (c, d) Scatter plots illustrated this gradient transformation, confirming that NPIs primarily act through DAN to maintain pre-intervention gradient architectures. TG2/CG2, post-intervention training/control group; DAN, dorsal attention network.

However, further analyses illustrated that removing the somato/motor network (SMN) and default mode network (DMN) also induced similar gradient alterations (Supplementary Fig. S2-4), implying these networks might also have the potential to maintain the cortical hierarchy. Given this, additional validation is required to determine whether only DAN substantially contributes to the observed improvements in global cognition associated with NPIs, or if it operates as part of an adaptive mechanism that is irrelevant to intervention effects.

### Gradient maintenance links to increases in functional connectivity strength within the dorsal attention network after intervention

Virtual lesion analyses identified the DAN as a core network maintaining gradient architecture after intervention. Given observed intervention-induced increases in within-FCS and DAN’s sensitivity to NPIs, we hypothesized that enhancing DAN’s internal functional integration (i.e., within-FCS increase in DAN) was the mechanism through which NPIs maintained gradient architecture in older adults. Based on the median within-FCS changes in DAN (Δwithin-FCS median = −0.0093), we divided TG2 participants (*n* = 112) into two subgroups (high/low-DAN, *n* = 56/56) and explored whether greater within-FCS enhancement in DAN after intervention was associated with maintained gradient organization.

We then computed post-intervention gradients separately for the high- and low-DAN subgroups from TG2 and compared them to the pre-intervention baseline TG1 pattern (*n* = 112) as well as the post-intervention natural aging CG2 pattern (*n* = 59). The high-DAN subgroup retained a gradient architecture highly consistent with TG1. Specifically, their primary and secondary gradients closely matched those of TG1 (G1-G1: *r* = 0.98, *p*_spin_ < 0.001; G2-G2: *r* = 0.99, *p*_spin_ < 0.001), with no cross-gradient similarity (G1-G2: *r* = 0.11, *p*_spin_ = 0.326; G2-G1: *r* = −0.10, *p*_spin_ = 0.652). In contrast, high-DAN gradients resembled CG2’s reversed pattern (G2-G1: *r* = 0.80, *p*_spin_ < 0.001; G1-G2: *r* = 0.79, *p*_spin_ = 0.004; Fig. 5a). Conversely, the low-DAN subgroup exhibited a flipped correspondence with TG1 (G1-G2: *r* = 0.99, *p*_spin_ < 0.001; G2-G1: *r* = 0.97, *p*_spin_ < 0.001), while closely aligning with the CG2 pattern (G1-G1: *r* = 0.80, *p*_spin_ = 0.004; G2-G2: *r* = 0.80, *p*_spin_ < 0.001; Fig. 5b). We also conducted parallel subgroup analyses for two other potential target networks (i.e., DMN and SMN) in TG2. While DMN showed a similar pattern to DAN, with gradient maintenance in the high-DMN group, SMN showed no such effect, implying it may reflect general adaptation in the elderly brain after intervention (Supplementary Fig. S5-9).

**Fig. 5.**
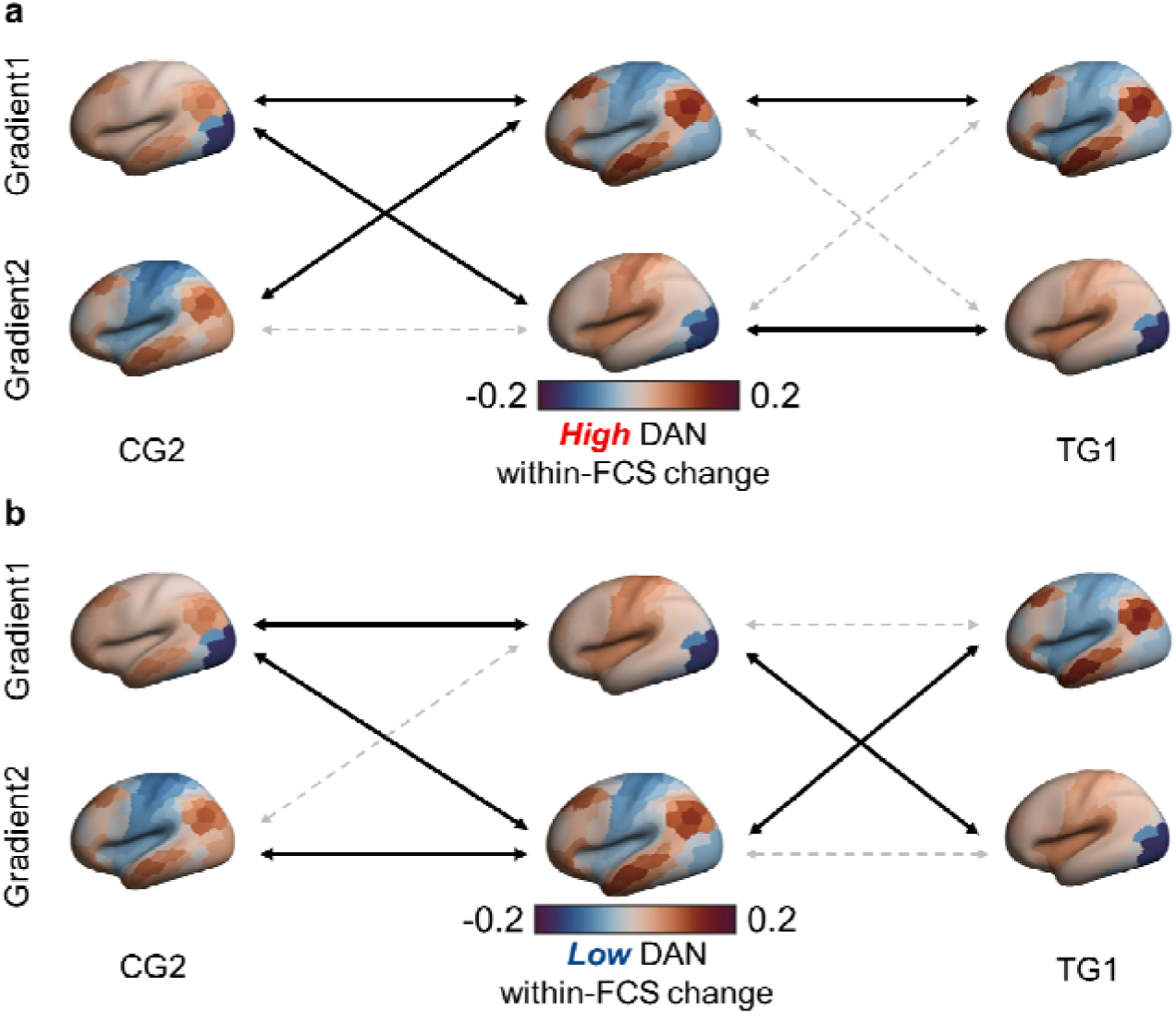
Gradient patterns in TG2 participants with higher and lower within-FCS changes in DAN after intervention. (a) In the high-DAN subgroup from TG2 (within-FCS in DAN increased above Δwithin-FCS median after NPIs), cortical gradient patterns closely resembled those of TG1, with no significant cross-gradient correspondence. In contrast, this subgroup diverged from the naturally aged pattern in CG2. (b) The low-DAN subgroup from TG2 (within-FCS in DAN increased below Δwithin-FCS median after NPIs) exhibited swapped gradients, with patterns similar to the CG2 and showing strong cross-gradient correlation with TG1 gradients. After intervention, TG2 participants with greater DAN integration enhancement maintained gradient architecture, whereas those with fewer increases followed an aging-related trajectory. Gradient colours (blue-red) represent gradient values. Black double arrows, significant positive associations (*p*_spin_ < 0.05); dashed gray arrows, null or negative associations. DAN, dorsal attention network; TG1, pre-intervention training group; CG2, post-intervention control group.

Together, these results provided an in vivo validation of the virtual lesion findings, confirming that the hierarchical brain organization in older adults can be effectively maintained after intervention by enhancing intra-network integration in DAN.

### Global cognition is related to functional connectivity strength within DAN, revealing dorsal attention network is the core convergent brain network across diverse NPIs attenuating age-related decline

Given the intervention-induced global cognition improvement (Fig. 1a) and gradient maintenance in TG caused by within-FCS increases in DAN (Fig. 5), our final investigation is whether the behavioral and brain intervention effects converge on DAN, the core shared brain network across diverse NPIs, as highlighted by prior analyses.

To examine the above hypothesis, we first compared MoCA score changes after intervention between high- and low-DAN subgroups (*n* = 49 and *n* = 48; Methods). As expected, the former showed significantly greater MoCA improvement than the latter (difference = 1.00, 95% CI = 0.00-1.00, *p* = 0.017, Cohen’s d = 0.277; Supplementary Fig. S7b, Mann-Whitney test due to non-normality). Similar comparisons based on DMN and SMN showed no significant differences (both *p* ≥ 0.664; Supplementary Fig. S8b-S9b), reinforcing DAN as the shared core network linking neural and cognitive benefits of NPIs in older adults.

We also constructed linear mixed-effect models predicting MoCA scores across all participants (*n* = 171), using within-FCS in DAN, DMN, SMN, or other networks as fixed effects separately, and repeated measures as random effects, with age, education, sex, and head motion as covariates (Methods). Results (Supplementary Table S3-5) revealed that DAN (*β* = 0.097, 95% CI = 0.026-0.168, *p* = 0.008) and DMN (*β* = 0.046, 95% CI = 0.004-0.089, *p* = 0.033) showed significant associations with MoCA, but only DAN survived Bonferroni correction (*p* = 0.023). SMN was not significantly related (*p* = 0.296) while other networks showed null results (all *p* ≥ 0.143; Supplementary Table S6-9).

We subsequently focused on brain and behavior changes after intervention only in TG (*n* = 112). A set of linear regression models examined whether changes in within-FCS (Δwithin-FCS) predicted MoCA improvement (Methods). Only Δwithin-FCS in DAN significantly predicted MoCA change (*β* = 0.101, 95% CI = 0.010-0.191, *p* = 0.031), with a trend towards significance after Bonferroni correction (*p* = 0.092). Neither DMN nor SMN showed significant effects (both *p* ≥ 0.283; Supplementary Table S10-12), while other networks showed null results (all *p* ≥ 0.561; Supplementary Table S13-16).

Finally, we assessed whether DAN in post-intervention TG (*n* = 112) remained predictive of global cognition. In linear models using post-intervention MoCA as the outcome (Methods), within-FCS in DAN (*β* = 0.180, 95% CI = 0.045-0.315, *p* = 0.010; corrected *p* = 0.030) and DMN (*β* = 0.107, 95% CI = 0.020-0.193, *p* = 0.018; corrected *p* = 0.054) remained strong predictors, while SMN remained non-significant (*p* = 0.315). Results from DAN, DMN, SMN, and four additional networks were listed (Supplementary Table S17-23).

Collectively, these findings indicated that intervention-enhanced within-FCS increase in DAN provided support for the maintenance of brain hierarchical organization in older adults and therefore facilitated global cognition, positioning DAN as the core convergent brain hub through which diverse NPIs achieve broad, far-transfer cognitive benefits attenuating age-related cognitive decline.

## Discussion

A pressing priority of heterogeneous non-pharmacological interventions in aging neuroscience is to reveal their universal brain pattern and underlying mechanism attenuating age-related cognitive decline, then furtherly identify the core convergent large-scale functional network central to their neuroplasticity in the elderly’s brain. In the current study, we constructed an NPI dataset based on four different types of intervention in older adults and investigated the intervention-general effects on both brain and behavior. Diverse NPIs help strengthen functional integration within the same network and maintain the original gradient pattern against aging, reflecting a significant intervention effect on brain hierarchical organization. Through virtual lesion analyses, we first uncovered that the dorsal attention network may be linked to the intervention-induced gradient maintenance in the cerebral macroscale organization. Next, we split the post-intervention training group into high- and low-subgroups according to the within-FCS changes median in DAN and found that elderly brain hierarchical organization can be effectively maintained by enhancing intra-network integration in DAN after intervention. Finally, the link between global cognition and within-FCS in DAN demonstrated that both the behavioral and brain intervention effects converge on DAN, once again indicating dorsal attention network is the core convergent hub across diverse non-pharmacological interventions attenuating age-related cognitive decline.

As intervention data used from our lab engaged multiple cognitive domains with indistinguishable contributions from four different intervention programs, we selected global cognition (MoCA scores) as the primary outcome. The results align with large, long-term, major intervention trials in older adults (FINGER, MYB) demonstrating NPI’s efficacy in reducing cognitive decline^29,30^, confirming our approach’s feasibility and effectiveness. Moreover, NPIs significantly altered brain functional connectivity. Older adults typically show reduced within-network and increased between-network functional connectivity compared to younger adults^11,12^, whose potential explanation for these age-related changes is dedifferentiation-the functional specialization loss in networks underpinned by more diffuse, nonspecific recruitment of brain regions underly cognitive decline^31^. In our study, NPIs induced reversible rsFC patterns with a significantly increased within-network functional connectivity and a decreased between-network functional connectivity trend in older adults, showing anti-aging changes with more interconnected brain systems. These findings reveal NPIs drive adaptive neuroplasticity through targeted intra-network integration, providing a mechanistic basis for delayed cognitive decline: brain regions become more specialized within functional networks like in adults counteracting age-related dedifferentiation.

Enhanced intra-network integration in older adults after intervention suggested a global shift of network architecture from dedifferentiation toward redifferentiation. To determine whether this change also reflects in more precise topographical organization, we examined macroscale connectivity gradients pre- and post-intervention, unveiling NPIs-induced aging reversing effect from an unexplored lens of gradient architecture. Pre-intervention gradients recapitulated established aging patterns^15,27^, showing the principal gradient from sensory/motor to transmodal association cortices and the secondary gradient from somatomotor to visual cortices. After NPIs, only the training group maintained original gradient organization while the control group exhibited somato/motor-visual and unimodal-transmodal as principal and secondary gradients (despite comparable explained variance with no significant difference; Supplementary Fig. S10). This illustrates that heterogeneous NPIs counteract age-related brain changes by maintaining a younger hierarchical gradient organization besides enhancing intra-network integration.

A fundamental gap in aging intervention research lies in the absence of a unified network-level framework through which diverse NPIs exert shared neuroplasticity enhancement, thereby attenuating age-related cognitive decline. Our findings address this gap by identifying enhanced intra-network functional integration within DAN as the core mechanism. Virtual lesion analyses in the post-intervention training group showed that a lesion simulation in DAN resembled normal aging gradient architecture in the post-intervention control group. Subsequent individual-level analysis in the same group further revealed that older adults with fewer DAN connections exhibited cortical organization similar to the post-intervention control group, whereas those with more DAN connections maintained the typical gradient architecture. Crucially, association analyses demonstrated that higher DAN connectivity reliably correlated with better global cognition across older adults. Together, these results highlight DAN’s pivotal role in attenuating age-related cognitive decline across heterogeneous NPIs. Diverse effective interventions significantly enhanced functional connectivity within DAN, thus maintaining brain functional gradient architecture while improving global cognition in older adults. Although our study combined four different NPIs, all inherently engaged attentional processes, a general though not unitary function where resource limits necessitate selection across perception, action, and memory^32^. Selective attention is essential for cognitive learning and might underlie its far transfer effects^33,34^, which is the ultimate goal of interventions aimed at cognitive enhancement in older people supported by DAN both from structural and functional levels^35–37^. DAN along with DMN represent two fundamental axes of brain functional organization, with the former controlling environmentally directed processes (e.g., perception and action) and the latter controlling internally directed processes^38,39^.

Neuroimaging studies have demonstrated that DAN maintained top-down modulatory signals for goal-relevant stimulus processing^40,41^.

Prior studies have emphasized attentional modulation on DAN connectivity as a key target for NPIs in older adults, regardless of intervention type. DAN plays a vital role in top-down control of sensory processing among different cognitive training^18^ and its changed connections related to far transfer^42^. Physical activity consistently benefits attention^43,44^, enhancing DAN interhemispheric connectivity to accelerate attention performance^45^. Multimodal intervention combining CI and PI also triggers DAN connectivity, where the increased functional connectivity strength in DAN positively correlated with task completion time^46^. Therefore, our findings support a model in which DAN plays a central convergent role across diverse NPIs in mediating behavioral outcomes. Given that cognitive decline in aging stems from disrupted coordination of large-scale systems conforming “last in, first out” model, including diminished within-network connectivity and aberrant between-network connectivity particularly in DAN and DMN^47–50^, along with an age difference in DAN gradients showing a functional connectivity profile more similar to that of other transmodal regions^27^, our results suggest that enhanced functional connectivity strength within dorsal attention network may serve as the neurofunctional biomarker of heterogeneous NPIs in maintaining hierarchical brain organization and attenuating age-related cognitive decline.

While present findings are promising, several limitations warrant consideration. First, though we revealed the core convergent network DAN underlying four different NPIs, our dataset could not encompass all intervention types (e.g., NIBS). Notably, prior work has provided partial support for selective effects of NIBS targeting dorsal and ventral attention networks during the early phases of multitask training^51^. Future studies should integrate broader NPI types in larger cohorts to validate generalizability. Second, participants in the control group were more self-selected with lower adherence than the intervention group. Despite homogenous baseline neuropsychological profiles and statistical corrections to minimize nonrandomized design effects, potential unmeasured biases such as socioeconomic level, daily enrichment, or motivation may have an impact on the results. Last, neuropsychological reassessment occurred without a washout period, risking practice effects. We minimize these effects on between-group differences via mixed-effect modeling of group × period interactions.

By characterizing macroscale cortex organization via functional connectome in older adults before and after interventions, we identify dorsal attention network as the core convergent brain hub through which diverse non-pharmacological interventions preserve gradient architecture and global cognition in aging. Specifically, older adults exhibiting increased functional connections within the dorsal attention network after intervention maintained typical gradient architecture and enhanced global cognition. These results advance current understanding of non-pharmacological interventions and aging-related neuroplasticity by uncovering a younger organization pattern and elucidating a shared neural mechanism underlying far-transfer effects of diverse NPIs, thus addressing a longstanding gap in the field through a unified network-level framework. While NPI-induced neuroplasticity involves multiscale interactions across environmental experience and biological systems, we establish DAN’s central role across diverse NPIs. All findings provide a methodological framework for the non-pharmacological interventions optimization in older adults, such as biomarker-guided targets for non-invasive brain stimulation. Collectively, these advances hold translational promise for promoting healthy aging, preventing cognitive decline, and enabling scalable implementation in clinical and public health contexts.

## Methods

### Participants

Data for this study were derived from four previously registered intervention trials in the Chinese Clinical Trial Registry (ChiCTR): Multimodal Intervention in Older Adults (http://www.chictr.org.cn, ChiCTR-PNRC-13003813); Combined Intervention of Aerobic Exercise and Video Game in Older Adults (http://www.chictr.org.cn, ChiCTR1900022702); Cognitive Training Modified Cognition in Older Adults (http://www.chictr.org.cn, ChiCTR-IOR-15006165); and Square Dance Enhanced Memory in Older Adults (http://www.chictr.org.cn, ChiCTR-IOR-17011668).

Participants who completed these trials within the intervention arms (including cognitive, exercise, dance, game, combined, and multimodal interventions) and had sufficient magnetic resonance imaging (MRI) quality were categorized as the Training Group (TG). Those from both passive and active control groups with adequate MRI quality were merged into the Control Group (CG). Details on participant recruitment and exclusion criteria across the four trials are presented in Fig. 1.

All participants provided written informed consent. These Trials were approved by the institutional review board of the Institute of Psychology, Chinese Academy of Sciences (H10025; H14020; H17008; H19006).

### Participant withdrawal

Participants who did not complete the entire trial, including those receiving the intervention but refusing the final resting-state functional MRI (rs-fMRI) scan, were classified as withdrawals. Their data were excluded from further analysis and kept confidential. Additionally, participants with excessive head motion artifacts in their rs-fMRI scans yielding insufficient time series data after scrubbing, were also classified as withdrawals. Their data were excluded and kept confidential, too.

Consequently, data from 171 participants who completed the intervention trials were included in the final analysis.

### Procedures

The study design, including the training and control group assignments, has been described in previous registrations and papers^52,53^. The overall procedure for participant recruitment, assignment, and intervention programs is illustrated in Fig. 1.

Inclusion criteria were as follows: (1) aged 53-80 years old; (2) ≥ 5 years of education; (3) normal global cognition; (4) right-handed; (5) no hearing, vision, or language deficits; (6) intact activities of daily living; (7) no psychiatric disorders, physical diseases, or neurological deficits affecting cognitive function; and (8) no contraindications for MRI.

Briefly, after providing informed consent and demographic information, all community-dwelling participants completed baseline assessments related to outcome measures, including neuropsychological tests (e.g., global cognition, executive function, and episodic memory). Only participants who voluntarily underwent an initial structural MRI and rs-fMRI scan and met the predefined inclusion criteria were formally enrolled in intervention trials.

Enrolled participants were assigned to either the training or control groups. The former received structured, non-pharmacological interventions delivered by trained volunteers, with durations ranging from 4 to 16 weeks. The specific interventions varied across trials and included standalone cognitive training, standalone physical exercise, and multimodal training (combined cognitive-physical training). Similarly, the latter engaged in trial-specific protocol-matched activities, including health lectures, maintenance of baseline lifestyle, or strenching exercise. The nature of both intervention and control conditions was clearly defined within each trial included in the present study. Upon finishing the assigned intervention or control sessions, participants were asked to complete the baseline-matched assessments, followed by a second MRI scan. Individuals failing to complete post-intervention measurements or provide high-quality data were excluded from final analyses. As a result, 112 training group participants and 59 control group participants were included in the final analysis as the TG and CG cohorts, respectively. The demographic information of all participants is listed in Supplementary Table S1.

All participants retained the right to withdraw from the study at any time for medical or personal reasons. Detailed assignment procedures and trial specifications are provided in Supplementary Table S1.

### MRI data acquisition

This study was conducted across multiple sites, with batch effects arising from the scanning of four intervention trials at different locations. MRI scans, including T1-weighted (T1w) and rs-fMRI, were obtained using 3-Tesla scanners with consistent parameters for each participant across both scans. Detailed information on the scanner types and protocols used for MRI across these intervention trials can be found in Supplementary Methods.

### MRI data preprocessing and postprocessing

Prior to processing, all data were organized and named according to the BIDS 1.6.0 specification. The anatomical T1w images were visually inspected to exclude individuals with substantial structural abnormalities, and the first five volumes of each rs-fMRI scan were discarded due to magnetic field equilibration.

After initial quality checks, T1w and rs-fMRI images were processed using the fMRIPrep (v23.2.1) and eXtensible Connectivity Pipeline-DCAN (XCP-D, v0.7.3) for surface-based analyses^54,55^. Both fMRIPrep and XCP-D are well-validated pipelines known for their robustness and reproducibility in fMRI processing. In brief, the preprocessing of rs-fMRI data included a series of steps: (1) generating skull-stripped reference volume; (2) estimating head motion parameters and head motion correction; (3) susceptibility distortion correction with fieldmap-less approach; (4) registration to high-resolution T1w images using boundary-based registration; (5) resampling the BOLD time series onto the fsaverage surface. Postprocessing was performed with XCP-D, using the 36p+Scrubbing model: (1) head radius = 50 mm for computing framewise displacement (FD), then scrubbing frames with FD > 0.5 mm; (2) band-pass filtering (0.01 Hz ≤ f ≤ 0.08 Hz); (3) removal of 36 nuisance factors (including six motion parameters, mean global signal, mean white matter signal, mean cerebrospinal fluid signal with their temporal derivatives, and quadratic expansion of six motion parameters, tissue signals and their temporal derivatives generated with fMRIPrep); (4) 6 mm spatial smoothing; (5) extraction of denoised time series using the Schaefer parcellations with 400 parcels^25^. In parallel, mean FD was calculated for each participant to account for residual motion effects. Participants with insufficient rs-fMRI scan volumes (< 240 seconds) were excluded from the final analysis.

### Functional Connectivity Strength

For each TG and CG participant, two 400 × 400 rsFC matrices (pre- and post-intervention) were generated. Each matrix was computed by Pearson’s *r* correlations between the time series of every pair of parcellations, then applying Fisher-z transformation to yield FCz values. Due to the multi-site nature of the study, involving different scanning parameters, the FCz matrices were harmonized using the neuroCombat toolbox v1.0.1^56^ before statistical analyses, with age, sex, and education as demographic covariates.

The functional connectivity strength (FCS) for each rs-fMRI scan was calculated at a network level. According to 7 networks from Yeo and colleagues^26^, all 400 cortical parcels were mapped into 7 networks. Within-FCS was calculated as the sum of FCz values between the parcellations within a given network, while between-FCS was calculated as the sum of FCz values between the parcellations of a given network and those outside of it. Additionally, longitudinal changes in FCS (ΔFCS) were computed by subtracting within- or between-FCS at initial scans from that at second scans, quantifying the effects of non-pharmacological interventions or normal aging on specific brain networks.

### Functional Connectivity Gradients

The functional connectivity gradients were calculated and displayed using BrainSpace v0.1.10^28^. First, for each trial period and group, FCz matrices were averaged across participants to create group-level matrices (TG1/CG1 for initial scans and TG2/CG2 for second scans). From four group-level FCz matrices, only the top 10% FCz values of each parcellation were retained, while negative and lower FCz values were zeroed to enforce sparsity, yielding an asymmetrical matrix. Then, the normalized angle distance was used to derive a symmetric affinity similarity matrix. Functional connectivity gradients were eventually calculated using diffusion embedding, a nonlinear dimensionality reduction technique. The gradient topography was visualized using colours from Scientific Colour Maps v8.0^57^.

### Group definition of high and low within-FCS in dorsal attention network

To examine the relationship between intervention-induced enhancements in global cognitive improvement and within-FCS in DAN, TG participants were stratified based on the cohort Δwithin-FCS (post-pre intervention) median. Participants exceeding this threshold were classified as the high-DAN subgroup, while participants below it were categorized as the low-DAN subgroup. Functional connectivity gradients were then recalculated for each subgroup, and changes in global cognition were compared between these two groups.

### Outcomes

Due to the heterogeneity of outcome measurements across four intervention trials, outcome selection was based on the most frequently assessed measures. All outcomes were evaluated longitudinally at pre- and post-intervention, with assessments independently conducted and double-checked by at least two trained volunteers.

Since the aim of this study was to reveal the intervention-general mechanisms across heterogeneous non-pharmacological interventions, we chose the global cognitive function, assessed using the Montreal Cognitive Assessment (MoCA), as the primary outcome. This neuropsychological test comprises seven cognitive domains (i.e., visuospatial, executive functioning, naming, attention, language, abstraction, delayed memory recall, and orientation), providing a comprehensive measure of intervention effects on global cognition.

The secondary outcomes related to cognition included measures of specific cognitive functions: verbal fluency, digit span-forward (DSF), digit span-backward (DSB), and Paired Associative Learning Test (PALT). These tests were used to assess distinct cognitive domains, covering language, executive function (working memory), and episodic memory.

### Statistical analysis

#### Intervention effects on cognition and functional connectivity strength

Intervention effects on cognition were assessed using 2 (group: TG vs. CG) × 2 (period: pre- vs. post-intervention) repeated-measures analysis of variance (ANOVA), with primary and secondary outcomes analyzed separately (see Supplementary Figures).

To evaluate intervention effects on functional connectivity strength, Δwithin-FCS was compared between TG and CG using the paired-sample Wilcoxon signed-rank test (due to non-normality), whereas Δbetween-FCS was analyzed using paired-sample t-tests.

#### Identification of the core convergent network

To investigate intervention-general networks, the principal and secondary functional connectivity gradients were derived for TG1, CG1, TG2, and CG2. The core convergent network role of DAN was tested by selectively dropping it (a simulation of lesion) from the TG2 FCz matrix and recomputing gradients. Pearson’s *r* was used to quantify the similarity between TG2 (DAN dropped) and CG2, with p-values obtained via two-sided spin tests. Similar analyses were also performed for other networks, identifying sensorimotor network (SMN) and default mode network (DMN) as additional possible networks (see Supplementary Figures).

#### Validation of global cognition-function connectivity strength links

To test the relationship between Δwithin-FCS in DAN and cognitive changes, TG2 participants were stratified as high- and low-DAN subgroups based on the Δwithin-FCS median. Gradient patterns of high- and low-DAN subgroups were separately compared to CG2 and TG1 using Pearson’s *r* (two-sided spin test), and cognitive outcome differences were assessed using independent-sample t-tests. Similar analyses were performed for SMN and DMN, two other probable networks.

#### Association analysis with global cognition (MoCA)

Three models were used to confirm the core convergent network role of DAN underlying diverse non-pharmacological interventions attenuating age-related cognitive decline rather than the other networks:

1. Global cognition in all participants (TG and CG). The association between within-FCS in DAN and global cognition was estimated with a linear mixed effect (LME) model. Within-FCS in DAN, together with age, education, sex, head motions (FD), and the intercept were set as the fixed effect factors. Subject (participant IDs), multi-measurements for a single participant were coded as an identical nominal variable, set as the random effect factor. The LME was conducted using the following formula:

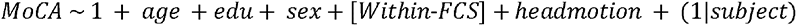

2. Changes in global cognition in TG participants. The association between Δwithin-FCS in DAN and changes in global cognition was estimated with a linear regression model (LM). Δwithin-FCS in DAN, together with age, education, sex, and the intercept were set as the predictors. The LM was conducted using the following formula:

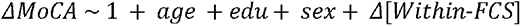

3. Global cognition in TG2 participants. The association between within-FCS in DAN and global cognition was estimated with an LM. Within-FCS in DAN, together with age, education, sex, head motions (FD), and the intercept were set as the predictors. The LM was conducted using the following formula:

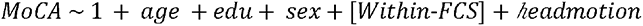

Parallel analyses were conducted for SMN and DMN (see Supplementary Tables). All models conducted Bonferroni corrections for multiple comparisons (across DAN, SMN, and DMN).

Statistical analyses were performed using BrainSpace v0.1.10 on MATLAB, SPSS v22.0, and R v4.2.1.

