## Supplementary for "Dorsal Attention Network as a Convergent Hub of Diverse Non-pharmacological Interventions Preserving Functional Gradients and Cognition in Aging"

**Summary of the content:**

I. Supplementary Methods
II. Supplementary Figures
III. Supplementary Tables

**I. Supplementary methods**

**MRI acquisition parameters in four different sites**

*Multimodal Intervention in Older Adults (site 1)*: A total of 16 participants of TG and 13 participants of CG were included in the analysis. All MRI data were obtained with a 3-Tesla Siemens Trio scanner located at the Beijing MRI Center for Brain Research. The rs-fMRI was collected using an EPI sequence with the following parameters: repetition time/echo time (TR/TE) = 2000 ms/30 ms; flip angle = 90°; field of view (FOV) = 200 mm × 200 mm; slice thickness = 3.0 mm; gap = 0.6 mm; acquisition matrix = 64 × 64; in-plane resolution = 3.125 × 3.125; 33 axial slices; 200 volumes (5 dummy scans). The structural imaging was collected using a MPRAGE T1-weighted sequence with the following parameters: TR/TE = 1900 ms/2.2 ms; acquisition matrix = 256 × 256; voxel size =1.0 × 1.0 × 1.0 mm^3^; flip angle = 9°; 176 slices.

*Combined Intervention of Aerobic Exercise and Video Game in Older Adults (site 2)*: A total of 65 participants of TG and 21 participants of CG were included in the analysis. All MRI data were obtained with a 3-Tesla GE MRI scanner (GE Discovery MR750) at the Magnetic Resonance Imaging Research Center, IPCAS. The rs-fMRI was collected using an EPI sequence with the following parameters: TR/TE = 2000 ms/30 ms; flip angle = 90◦; field of view = 224 × 224 mm; acquisition matrix = 64 × 64; slice thickness = 3.5 mm; voxel size = 3.5 × 3.5 × 3.5 mm^3^; 37 slices; 240 volumes (5 dummy scans). The structural imaging was collected using a 3D T1-weighted sequence with the following parameters: TR/TE = 6.7 ms/2.9 ms; inversion time = 450 ms; acquisition matrix = 256 × 256; voxel size =1.0 × 1.0 × 1.0 mm^3^; flip angle = 12°; 192 slices.

*Cognitive Training Modified Cognition in Older Adults (site 3)*: A total of 15 participants of TG and 13 participants of CG were included in the analysis. All MRI data were obtained with a 3-Tesla Siemens Trio scanner located at the Beijing MRI Center for Brain Research. The rs-fMRI was collected using an EPI sequence with the following parameters: repetition time/echo time (TR/TE) = 2000 ms/30 ms; flip angle = 90°; field of view (FOV) = 200 mm × 200 mm; slice thickness = 3.0 mm; gap = 0.6 mm; acquisition matrix = 64 × 64; in-plane resolution = 3.125 × 3.125; 33 axial slices; 200 volumes (5 dummy scans). The structural imaging was collected using a MPRAGE T1-weighted sequence with the following parameters: TR/TE = 1900 ms/2.2 ms; acquisition matrix = 256 × 256; voxel size =1.0 × 1.0 × 1.0 mm^3^; flip angle = 9°; 176 slices.

*Square Dance Enhanced Memory in Older Adults (site 4)*: A total of 16 participants of TG and 12 participants of CG were included in the analysis. All MRI data were obtained with a 3-Tesla Siemens MAGNETOM Prisma scanner located at the Center for MRI Research, Peking University. The rs-fMRI was collected using an EPI sequence with the following parameters: repetition time/echo time (TR/TE) = 2000 ms/30 ms; flip angle = 90°; field of view (FOV) = 224 mm × 224 mm; slice thickness = 2.0 mm; voxel size = 2.0 × 2.0 × 2.0 mm^3^; 62 slices; 240 volumes (5 dummy scans). The structural imaging was collected using a MPRAGE T1-weighted sequence with the following parameters: TR/TE = 2400 ms/2.2 ms; FOV = 256 × 256 mm^2^; voxel size =0.75 × 0.75 × 0.75 mm^3^; flip angle = 8°; 224 slices.

**II. Supplementary figures**

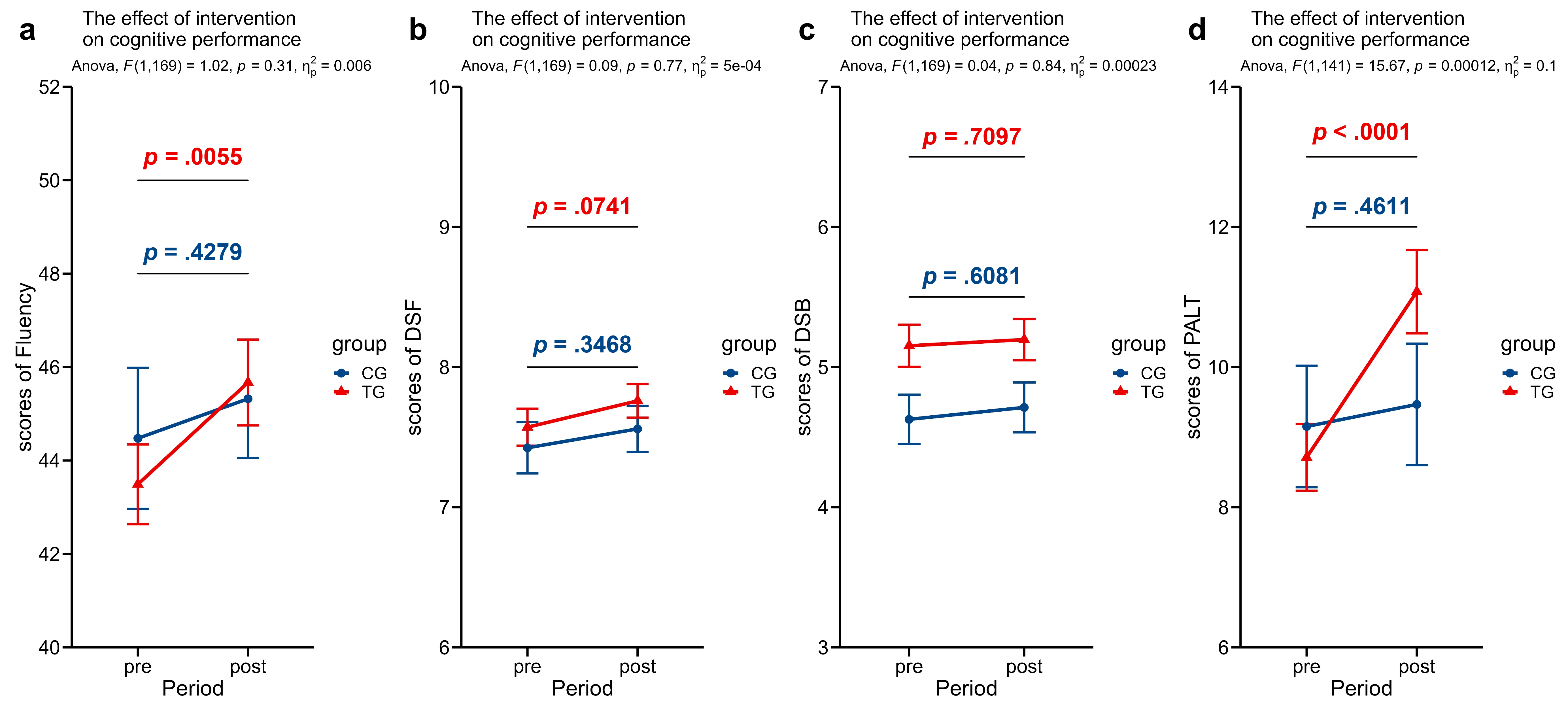

**Fig. S1. Intervention effects on secondary outcomes.** No significant interaction effect of group (TG vs. CG) × period (pre- vs. post-intervention) was observed for verbal fluency **(a)**, DSF **(b)**, or DSB **(c)** (all *ps* > 0.310), suggesting limited NPIs effects on these specific measures. However, simple main effects indicated a potential training-related enhancement in verbal fluency within the TG (*p* = 0.006). Conversely, the PALT **(d)** showed a significant interaction (*p* < 0.001), driven by improved performance in the TG (*p* < 0.001) with no significant change in the CG (*p* = 0.461). These findings demonstrate that NPIs exert intervention effects on multiple domains, supporting the existence of far-transfer effects. Higher scores indicate better performance. Error bars represent the standard error of the means. TG/CG, Training Group/Control Group; DSF/DSB, Digit Span Forward/Backward; PALT, Paired Associative Learning Test; NPIs, Non-pharmacological Interventions.

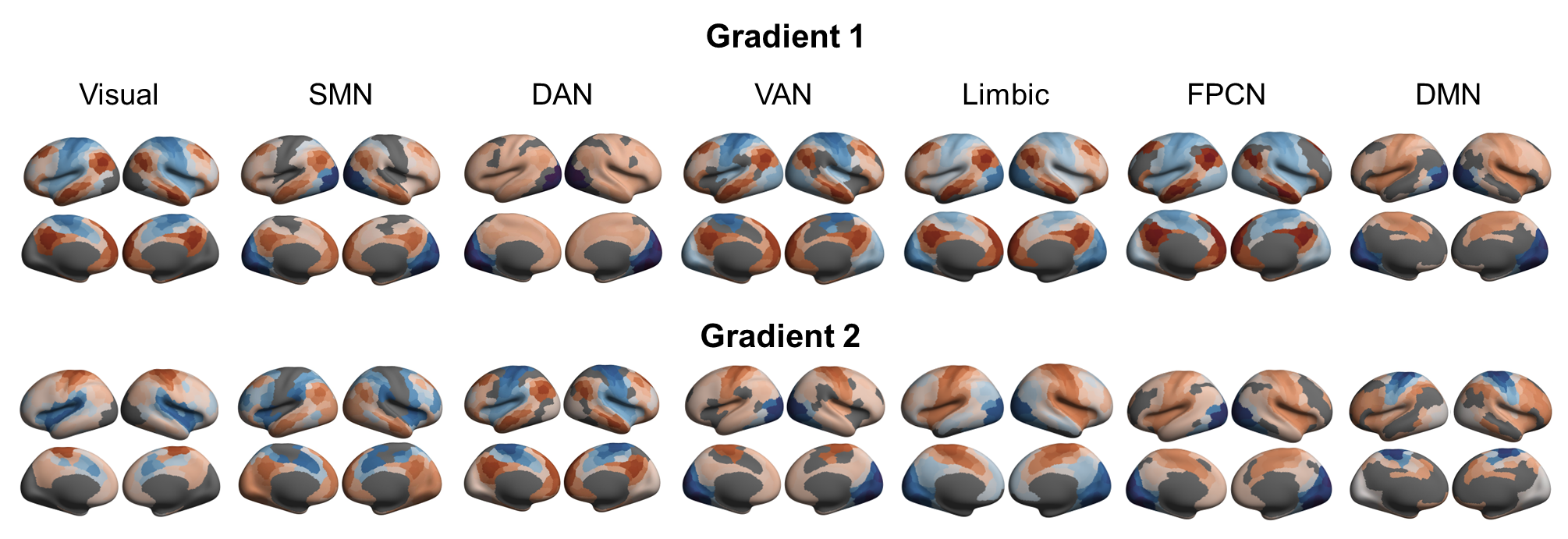

**Fig. S2. Gradient maps with functional networks lesioned separately.** Systematic virtual lesion analyses revealed distinct preservation patterns in the TG2: When somatomotor network (SMN), default mode network (DMN), or dorsal attention network (DAN) were removed, the gradient patterns became consistent with the CG2 pattern. Conversely, the dropout of other networks generated transmodal organization in the primary gradient without reversal, maintaining the original pattern of TG2. Nevertheless, interpretation requires caution given inherent biases from variations in network sizes across the cortical sheet. TG/CG, Training Group/Control Group.

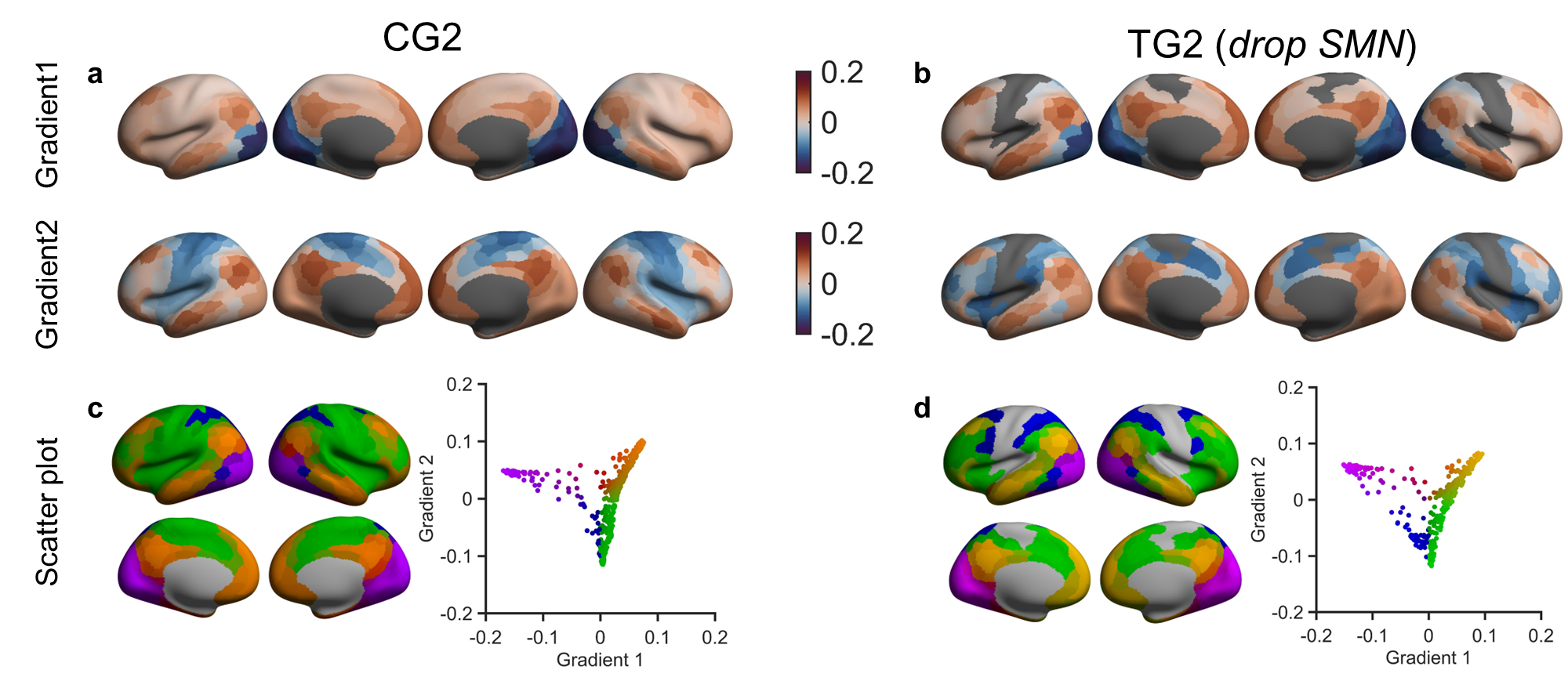

**Fig. S3. The Somato/motor network may also prevent gradient reversal.** The primary and secondary gradients of CG2 **(a)**, and SMN-dropped TG2 **(b)**. The primary and secondary gradients of TG2 (drop SMN) also closely matched those of CG2 (G1-G1: *r* = 0.98, *p*_spin_ < 0.001; G2-G2: *r* = 0.93, *p*_spin_ < 0.001; 1000 permutations), whereas no such similarity was found between TG2’s primary gradient and CG2’s secondary gradient, or vice versa (*ps* ≥ 0.196). Scatter plots **(c, d)** illustrate this gradient transformation.

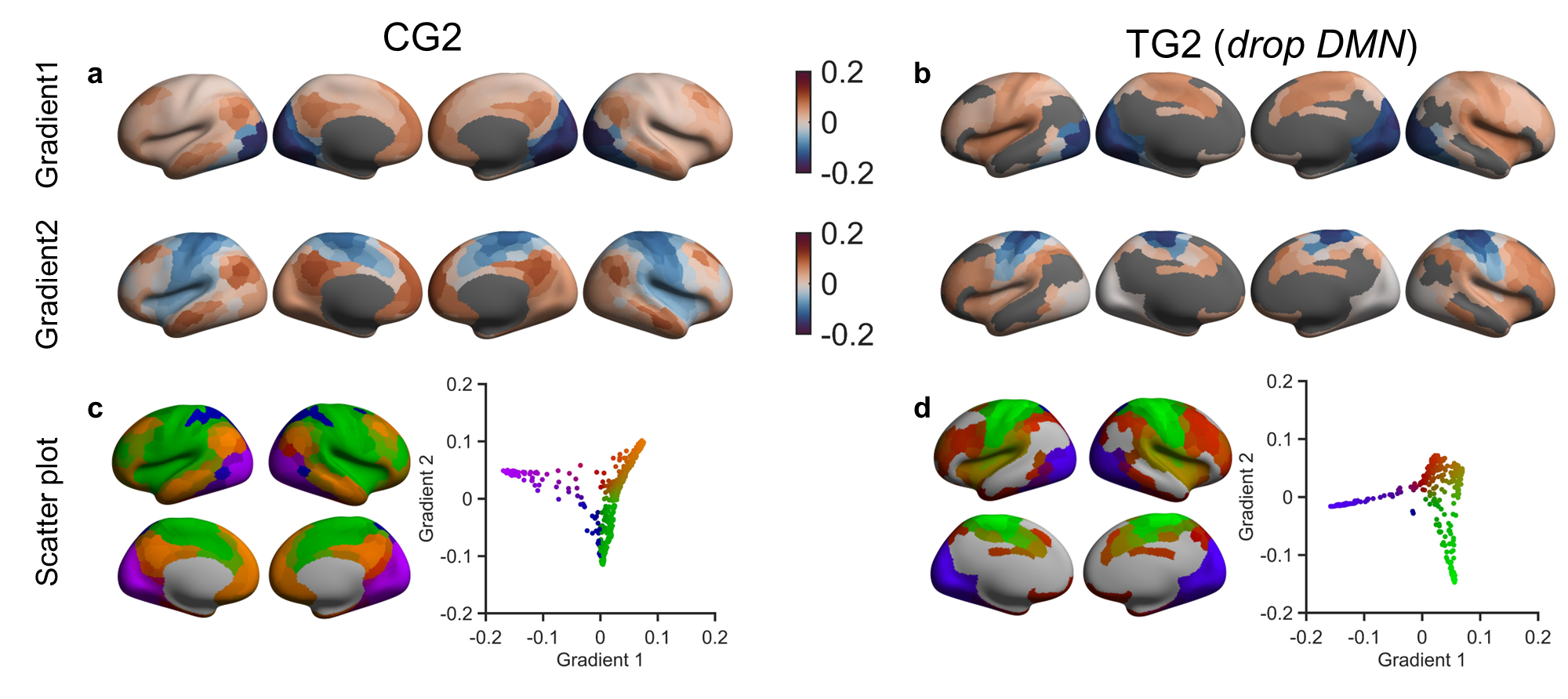

**Fig. S4. The Default mode network may also prevent gradient reversal.** The primary and secondary gradients of CG2 **(a)**, and DMN-dropped TG2 **(b)**. The primary and secondary gradients of TG2 (drop DMN) also closely matched those of CG2 (G1-G1: *r* = 0.92, *p*_spin_ < 0.001; G2-G2: *r* = 0.56, *p*_spin_ = 0.019; 1000 permutations), whereas no such similarity was found between TG2’s primary gradient and CG2’s secondary gradient, or vice versa (*ps* ≥ 0.218). Scatter plots **(c, d)** illustrate this gradient transformation.

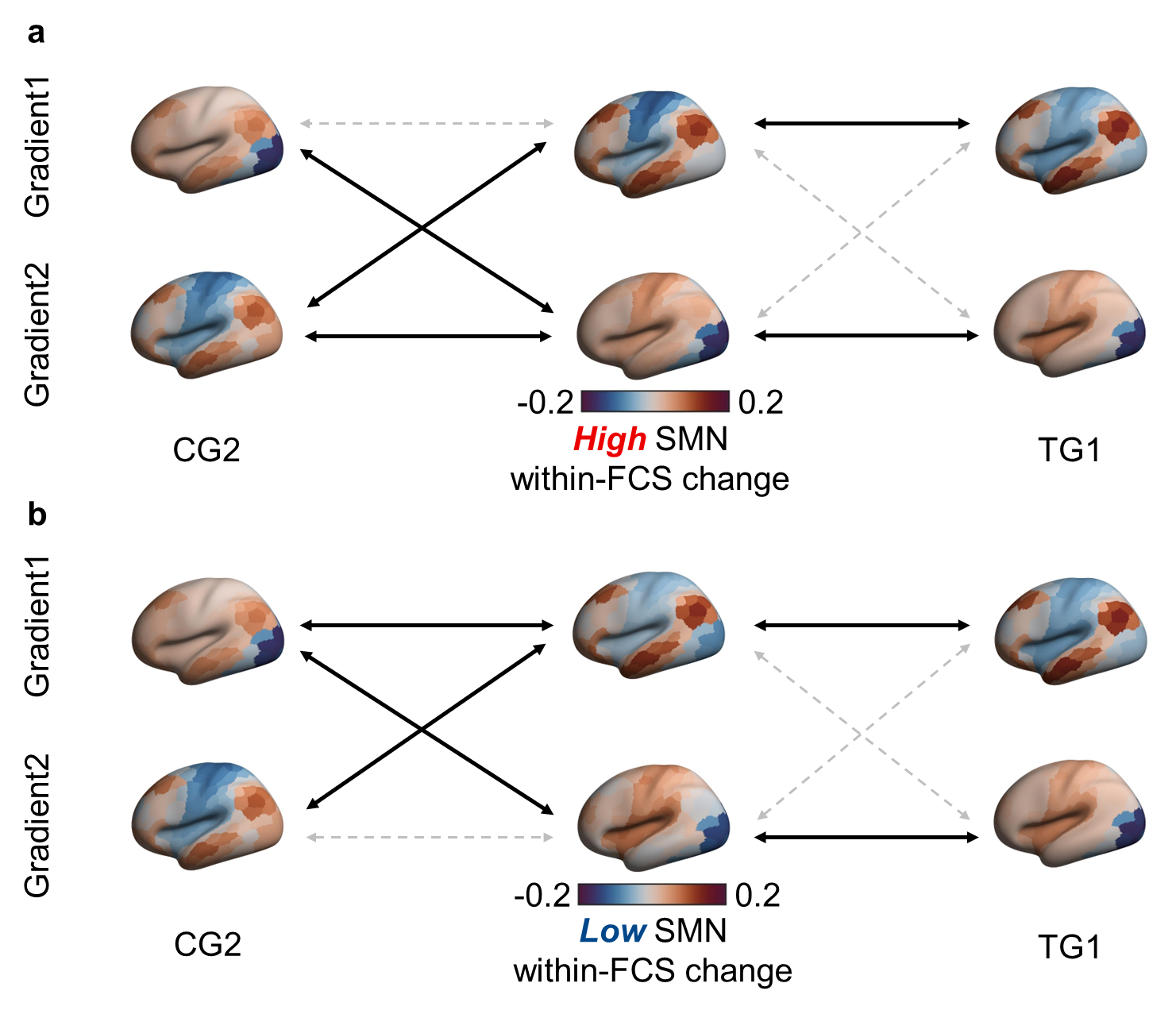

**Fig. S5. Gradient patterns in TG2 participants with higher and lower intervention-induced increases in SMN within-FCS.** **(a)** In the high-SMN subgroup (with greater within-FCS increase in SMN after intervention), cortical gradient patterns remained consistent with the TG prior to intervention (TG1; G1-G1: *r* = 0.94, *p*_spin_ < 0.001; G2-G2: *r* = 0.98, *p*_spin_ < 0.001), and showed divergence from the CG post-intervention pattern (CG2; G1-G2: *r* = 0.90, *p*_spin_ < 0.001; G2-G1: *r* = 0.91, *p*_spin_ < 0.001). **(b)** Similarly, the low-SMN subgroup also maintained a TG1-like gradient architecture (G1-G1: *r* = 0.96, *p*_spin_ < 0.001; G2-G2: *r* = 0.97, *p*_spin_ < 0.001) and differed from CG2 (G1-G2: *r* = 0.69, *p*_spin_ < 0.001; G2-G1: *r* = 0.70, *p*_spin_ = 0.011). Unlike DAN, the preservation of gradient architecture in SMN is not modulated by the effect size of intervention, but may instead reflect a general adaptive mechanism after NPIs. Gradient colours (blue-red) represent gradient values. Black double arrows, significant positive correlations (*p*_spin_ < 0.05); dashed gray arrows, non-significant or negative associations. TG/CG, Training/Control Group; SMN, somato/motor network. The scatter plots are displayed in Supplementary Fig. S7a, using TG1 (Supplementary Fig. S7b) as the template.

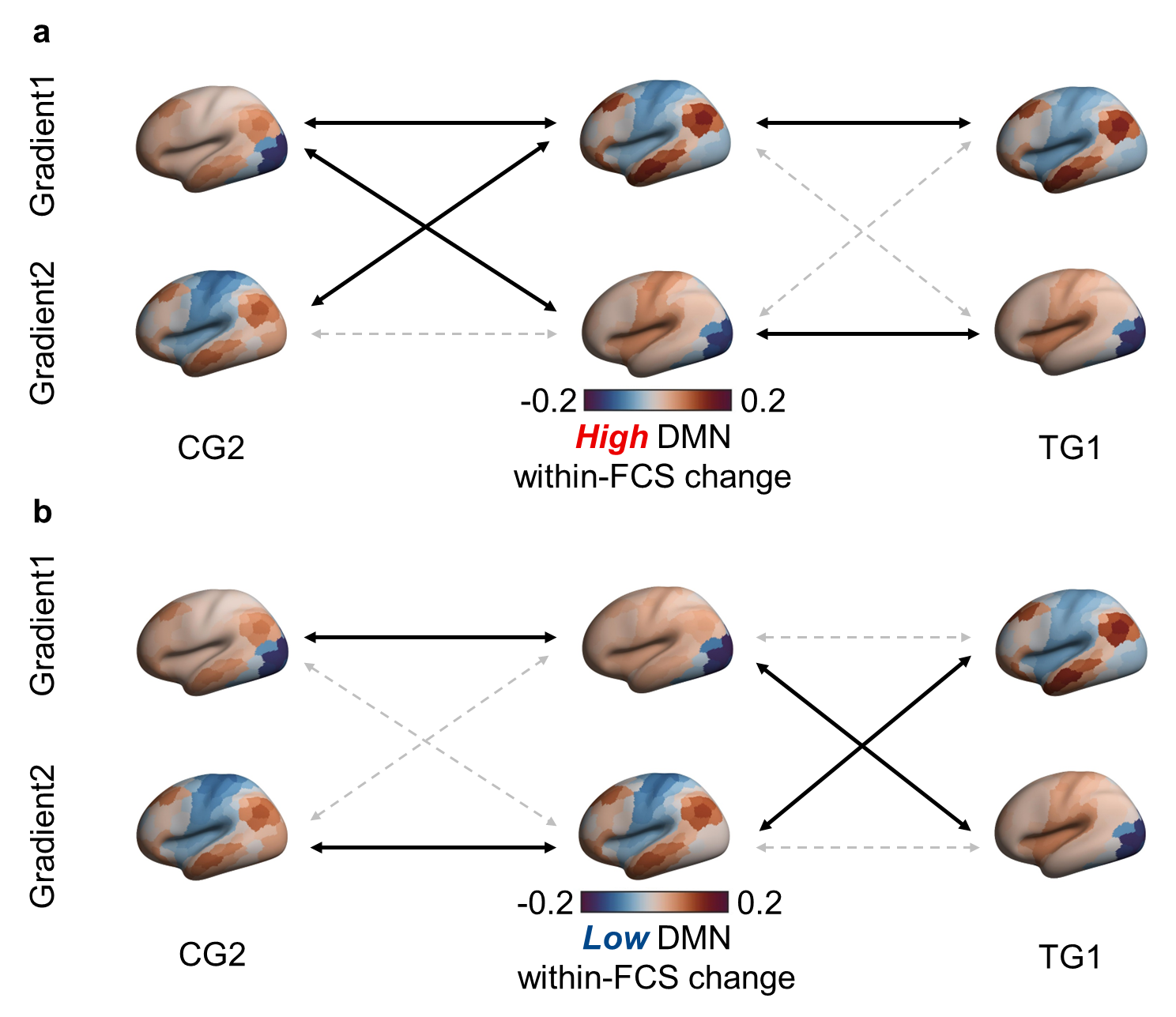

**Fig. S6. Gradient patterns in TG2 participants with higher and lower intervention-induced increases in DMN within-FCS.** **(a)** In the high-DMN subgroup, cortical gradient patterns closely resembled TG1 (G1-G1: *r* = 0.99, *p*_spin_ < 0.001; G2-G2: *r* = 0.99, *p*_spin_ < 0.001). This subgroup diverged from CG2 participants (G1-G2: *r* = 0.84, *p*_spin_ < 0.001; G2-G1: *r* = 0.84, *p*_spin_ < 0.001). **(b)** However, the low-DMN subgroup exhibited gradient reversal, with gradients matching CG2 (G1-G1: *r* = 0.95, *p*_spin_ < 0.001; G2-G2: *r* = 0.95, *p*_spin_ < 0.001) and cross-gradient similarity with TG1 (G1-G2: *r* = 0.97, *p*_spin_ < 0.001; G2-G1: *r* = 0.93, *p*_spin_ < 0.001). Gradient colours (blue-red) represent gradient values. Black double arrows, significant positive associations (*p*_spin_ < 0.05); dashed gray arrows, null or negative associations. TG/CG, Training/Control Group; DMN, default mode network. The scatter plots are displayed in Supplementary Fig. S8a, using TG1 (Supplementary Fig. S8b) as the template.

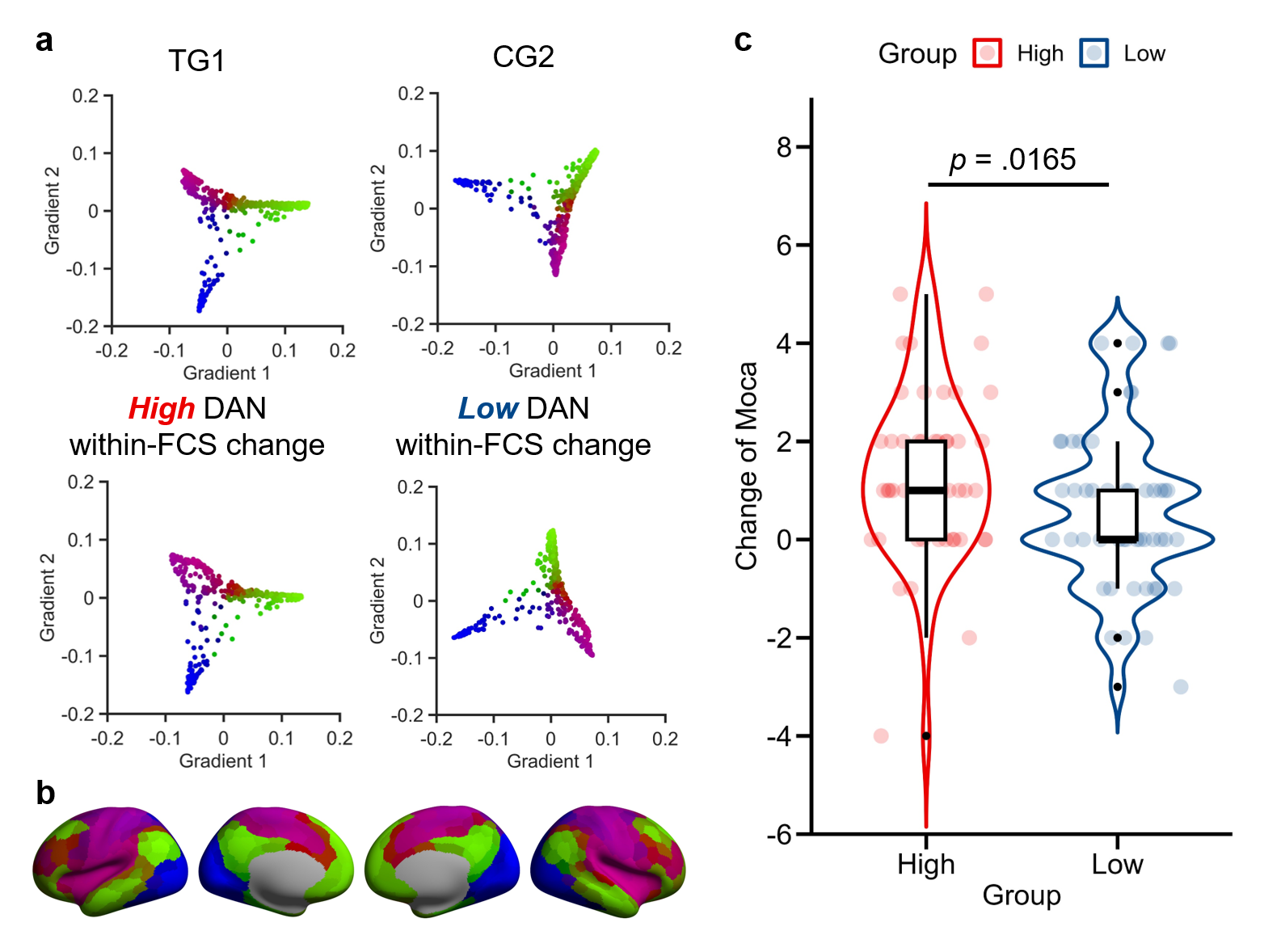

**Fig. S7. Intervention-induced within-FCS increases in DAN support preserved gradient architecture and cognitive improvements.** **(a)** Scatter plots show gradient values of the Schaefer parcellations with 400 cortical parcels. High-DAN subgroup from TG2 (within-FCS in DAN increased above median Δwithin-FCS after NPIs) retained the TG1-like gradient architecture, while low-DAN subgroup from TG2 (within-FCS in DAN increased below median Δwithin-FCS after NPIs) and CG2 both showed reversed gradients, suggesting disrupted hierarchy organization. **(b)** Gradient template from TG1 participants. **(c)** MoCA score changes after the intervention in High- and low-DAN subgroups. The former showed greater global cognition improvement than the latter (Mann-Whitney test; difference = 1.00, 95% CI = 0.00-1.00, *p* = 0.017, Cohen’s d = 0.277).

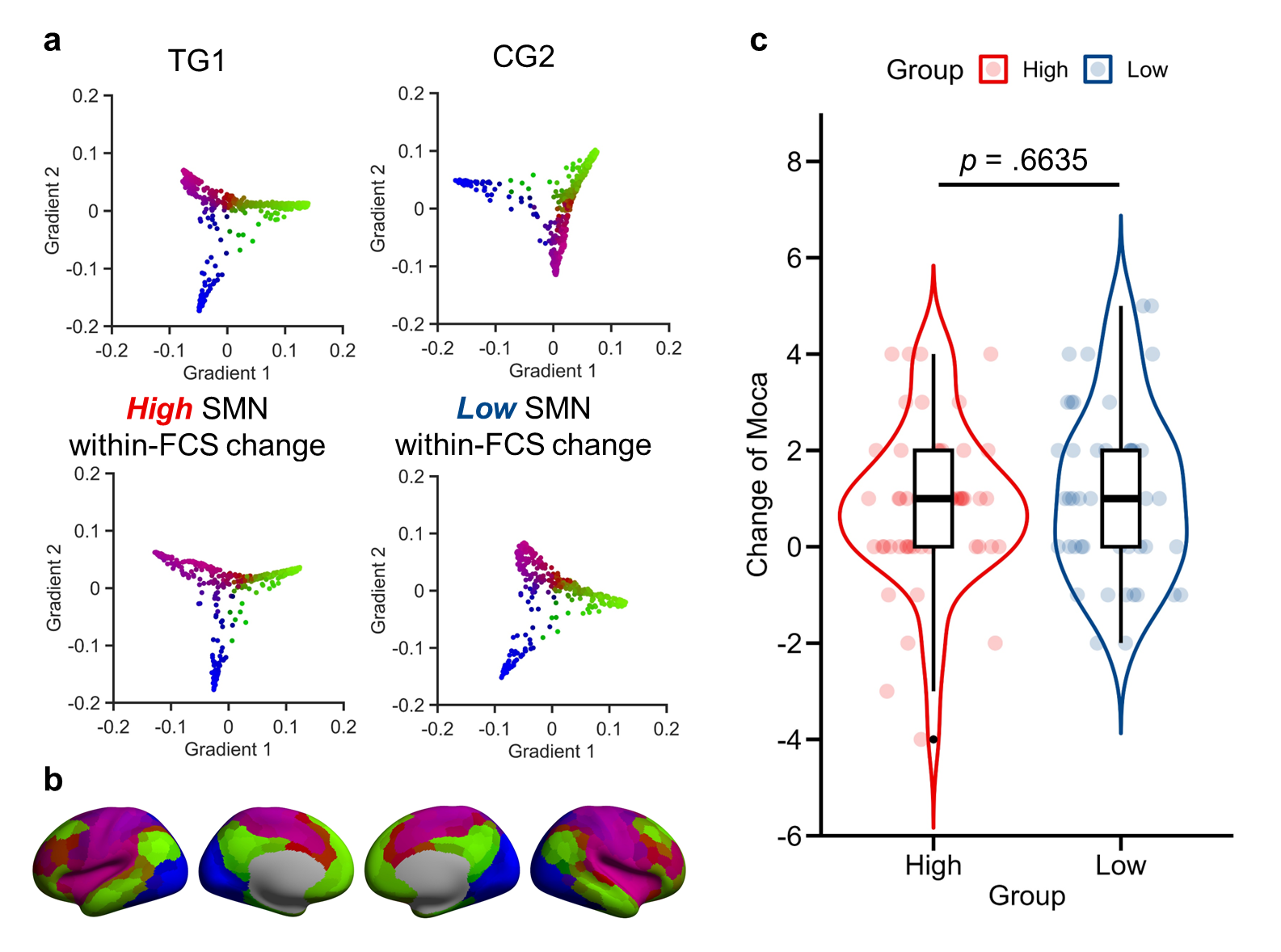

**Fig. S8. Intervention-induced within-FCS increase in SMN maintains gradients but lacks cognitive impact. (a)** Scatter plots show gradient values of 400 cortical parcels using TG1 as the reference. Both high- and low-SMN subgroups retained the TG1-like gradient architecture, despite the low-SMN subgroup showing a slight spin. **(b)** Gradient template from TG1 participants. **(c)** MoCA score changes before and after intervention (high: *n* = 51; low: *n* = 46). No significant difference was found between the two subgroups (Mann-Whitney test; difference = 0.00, 95% CI = -1.00-1.00, *p* = 0.664, Cohen’s d = 0.051).

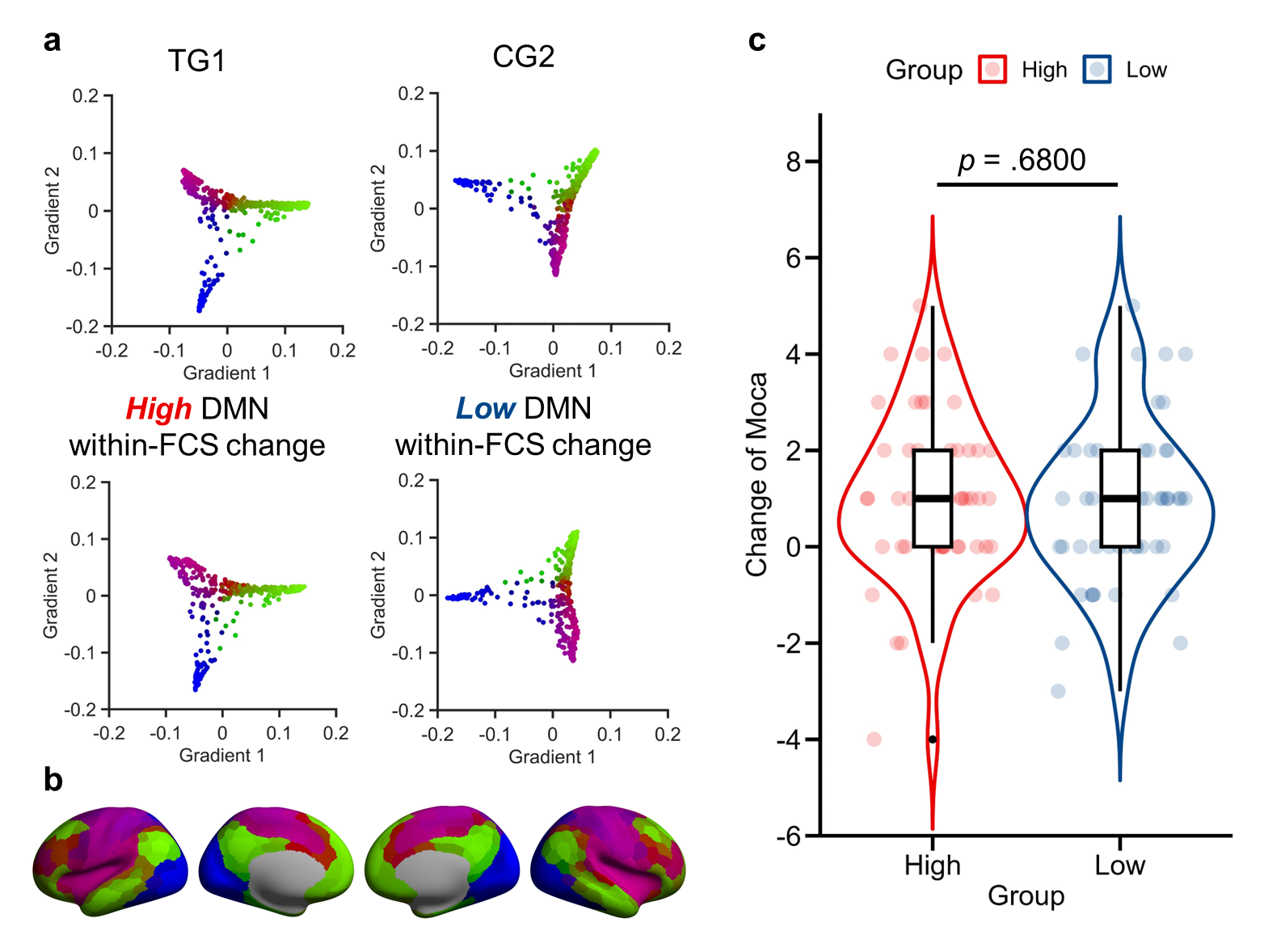

**Fig. S9. Intervention-induced within-FCS increase in DMN maintains gradients but lacks cognitive impact. (a)** Scatter plots show gradient values of 400 cortical parcels using TG1 as the reference. High-DMN subgroup retained the TG1-like gradient architecture, while low-DMN and CG2 showed reversed gradients. **(b)** Gradient template from TG1 participants. **(c)** MoCA score changes before and after intervention (high: *n* = 48; low: *n* = 49). No significant difference was found between the two subgroups (independent samples *t*-test; difference = -0.14, 95% CI = -0.83-0.54, *p* = 0.680, Cohen’s d = -0.084).

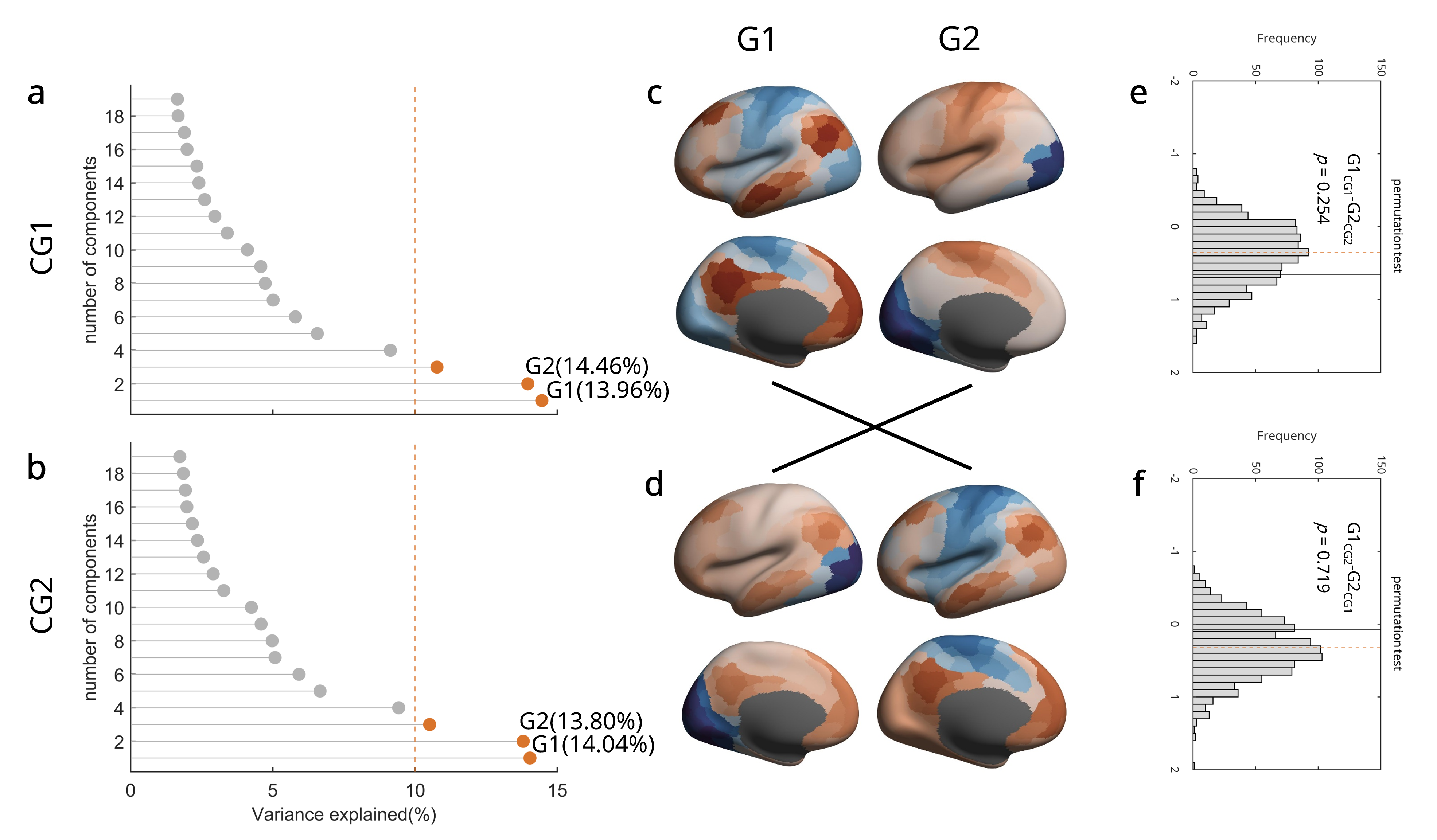

**Fig. S10. The principal and secondary gradients reversed in the control group with no variance significance.** Permutation analyses were conducted to examine the significance of the gradient reversal in CG participants. The 118 FCz matrices (59 for pre-intervention and 59 for post-intervention) of CG participants were assigned randomly to two datasets corresponding to the number of matrices (pre-/post-intervention: 59/59) for 1000 permutations. The gradient maps were then recomputed and the explained variances of the first two gradients were extracted to generate a set of null models. The original explained variance of CG1 (**a**, G1: 0.1446; G2: 0.1395) and CG2 (**b**, G1: 0.1404; G2: 0.1380), as well as corresponding gradient architectures (**c**, CG1; **d**, CG2), were displayed. The permutation analyses demonstrate that no significant decrease in the amount of variance accounted for by the association cortex anchored gradient (pre-post: 0.0065) was found in the CG participants (*p* = 0.254, 1000 permutations; **e**). Consistently, the amount of variance accounted for by the unimodal anchored gradient also shows no significant increase (post-pre: 0.0007) in the CG participants (*p* = 0.719, 1000 permutations; **f**). CG, Control Group.

**III. Supplementary tables**

**Supplementary Table S1.** Detailed allocation procedures and trial specifications of the four intervention trials analyzed in the present study.

| **Previous trials** | **Multimodal Intervention** | | **Combined Intervention** | | **Cognitive Training** | | **Square Dance** | |
| --- | --- | --- | --- | --- | --- | --- | --- | --- |
| **Study type** | Interventional study | | Interventional study | | Interventional study | | Interventional study | |
| **Study design** | Nonrandomized Intervention Trial | | Nonrandomized Intervention Trial | | Randomized Parallel Controlled Trial | | Randomized Parallel Controlled Trial | |
| **Allocation** | Participants from two similar communities were blind to the study design and the two communities were randomly allocated to the intervention or control group. | | Participants were blind to the aims and hypotheses of the study and randomly allocated due to their self-preferences and time conflicts. | | Participants were blind to the study design and randomly allocated to the intervention or control group. | | A single-blind RCT. Participants were blind to the study design and randomly allocated to the intervention or control group (1:1) by an independent volunteer using the randomisation programme online generation tool (http://www.randomization.com) | |
| **Intervention program** | ***Intervention*** | ***Passive control*** | ***Intervention*** | ***Passive control*** | ***Intervention*** | ***Passive control*** | ***Intervention*** | ***Active control*** |
|  | 6-week multimodal activities, including cognitive intervention (3 sessions/week, 1h/session), Tai Chi exercise (3 sessions/week, 1h/session), group counseling (1 session/week, 1.5h/session) | 2 times of 2h health lecture | 16-week combined training: 2 sessions/week, 1h/session  16-week aerobic cycling: 2 sessions/week, 1h/session  16-week video game: 2 sessions/week, 1h/session | Lifestyles remained unchanged. | 4-week cognitive training, including memory strategy training and executive function training (3 sessions/week, 1.5h/session). Participants were assigned homework after  each training session to practice at home. | 4-week lectures on health and aging (3 sessions/week, 1.5h/session) | 14-week square dance (moderate physical activity): 2 sessions/week, 1.5h/session | 14-week stretch training: 2 sessions/week, 1.5h/session |

**Supplementary Table S2.** Basic characteristics and average head motion during each scan in four intervention trials (N = 171).

|  | **Multimodal Intervention** | | **Combined Intervention** | | **Cognitive Training** | | **Square Dance** | |
| --- | --- | --- | --- | --- | --- | --- | --- | --- |
| **Characteristics** | TG (*n* = 16) | CG (*n* = 13) | TG (*n* = 65) | CG (*n* = 21) | TG (*n* = 16) | CG (*n* = 12) | TG (*n* = 15) | CG (*n* = 13) |
| **Demographics** |  |  |  |  |  |  |  |  |
| Age (year) | 68.6(5.7) | 71.9(4.3) | 63.1(4.6) | 63.3(5.0) | 74.5(3.5) | 71.9(5.8) | 63.0(5.7) | 62.9(4.5) |
| Females | 7(43.8) | 5(38.5) | 52(80.0) | 18(85.7) | 12(75.0) | 8(66.7) | 14(93.3) | 13(100.0) |
| Education | 13.3(3.9) | 14.8(3.3) | 11.5(2.1) | 11.6(2.7) | 15.6(2.4) | 15.0(3.4) | 12.7(2.4) | 11.0(3.3) |
| **Global cognition** |  |  |  |  |  |  |  |  |
| MoCA (pre) | 26.2(2.7) | 25.2(2.6) | 26.1(2.2) | 25.7(2.0) | 26.1(2.0) | 27.3(1.4) | / | / |
| MoCA (post) | 27.3(2.8) | 26.4(2.8) | 27.0(2.3) | 25.7(2.0) | 27.1(1.7) | 27.2(17) | / | / |
| **Single-domain cognition** |  |  |  |  |  |  |  |  |
| Fluency (pre) | 43.5(9.8) | 41.2(14.8) | 42.8(8.3) | 45.4(11.9) | 45.1(11.5) | 44.6(10.3) | 44.7(8.7) | 46.2(8.8) |
| Fluency (post) | 49.2(11.0) | 45.9(8.7) | 44.6(8.5) | 43.2(10.0) | 48.6(11.7) | 47.9(8.6) | 43.4(10.1) | 45.8(11.5) |
| DSF (pre) | 7.5(1.3) | 7.4(1.3) | 7.6(1.4) | 7.3(1.6) | 7.1(1.4) | 7.3(1.3) | 8.1(1.5) | 7.8(1.4) |
| DSF (post) | 7.6(1.6) | 7.4(1.3) | 7.8(1.2) | 7.4(1.2) | 7.6(1.2) | 7.4(1.1) | 7.9(1.2) | 8.1(1.4) |
| DSB (pre) | 5.4(1.9) | 4.1(1.0) | 5.2(1.6) | 4.7(1.2) | 4.8(1.4) | 5.0(1.5) | 4.9(1.4) | 4.8(1.7) |
| DSB (post) | 5.3(1.9) | 4.6(1.4) | 5.3(1.6) | 4.5(1.0) | 4.9(1.3) | 4.8(1.5) | 5.0(1.1) | 5.0(1.7) |
| PALT (pre) | 2.4(1.0) | 2.3(1.5) | 9.0(3.9) | 10.6(3.1) | 13.8(2.4) | 14.0(6.1) | / | / |
| PALT (post) | 3.4(1.7) | 2.5(1.0) | 11.4(4.6) | 10.7(3.0) | 17.6(4.2) | 14.9(5.6) | / | / |
| **Emotion** |  |  |  |  |  |  |  |  |
| SAS (pre) | / | / | 29.9(4.3) | 31.4(6.9) | 28.6(5.8) | 29.1(5.3) | 25.9(3.7) | 26.4(6.2) |
| SAS (post) | / | / | 30.9(5.6) | 31.4(6.3) | 25.4(3.3) | 26.6(6.5) | 27.0(4.9) | 26.6(6.6) |
| CES-D (pre) | 7.6(7.2) | 6.4(6.7) | 4.3(6.4) | 8.2(8.3) | 6.3(7.4) | 7.3(8.3) | 7.0(4.5) | 6.8(6.7) |
| CES-D (post) | 5.1(3.6) | 4.8(4.8) | 5.0(6.5) | 6.6(6.9) | 7.0(7.4) | 5.2(5.6) | 6.1(4.4) | 6.1(6.3) |
| **Head motion** |  |  |  |  |  |  |  |  |
| FD1 (initial scan) | 0.17(0.05) | 0.19(0.07) | 0.11(0.05) | 0.13(0.04) | 0.20(0.06) | 0.22(0.06) | 0.16(0.07) | 0.14(0.04) |
| FD2 (final scan) | 0.18(0.06) | 0.19(0.07) | 0.12(0.05) | 0.14(0.05) | 0.20(0.05) | 0.21(0.07) | 0.16(0.07) | 0.14(0.04) |

*Note:* data are n (%) or mean (s.d.). MoCA, Montreal Cognitive Assessment; Fluency, verbal fluency; DSF/DSB, digital span-forward/backward; PALT, Paired Associative Learning Test; SAS, self-rating anxiety scale; CES-D, Center for Epidemiological Studies-Depression; pre/post, pre-intervention/post-intervention.

$$Cognitive Score\approx1 + age +edu+ sex+[Within\text{-}FCS]+headmotion + (1|subject)$$

**In all TG and CG participants dataset (*n* = 171):**

**Supplementary Table S3.** Linear mixed effect models assessing relationships between the within-FCS of the somato/motor network (SMN) and MoCA

| IV: MoCA | Estimate *β* | SE | t | p |
| --- | --- | --- | --- | --- |
| Intercept | 26.86438 | 1.92637 | 13.946 | <0.0001 |
| Age | -0.02786 | 0.0288 | -0.967 | 0.3349 |
| Sex | -0.60987 | 0.38355 | -1.59 | 0.1141 |
| Edu | 0.14226 | 0.05763 | 2.469 | 0.0148 |
| FD | -1.91366 | 2.26153 | -0.846 | 0.3982 |
| SMN | 0.01871 | 0.01785 | 1.048 | 0.2955 |

**Supplementary Table S4.** Linear mixed effect models assessing relationships between the within-FCS of the dorsal attention network (DAN) and MoCA

| IV: MoCA | Estimate *β* | SE | t | p |
| --- | --- | --- | --- | --- |
| Intercept | 25.89076 | 1.93658 | 13.369 | <0.0001 |
| Age | -0.02265 | 0.02861 | -0.791 | 0.42999 |
| Sex | -0.58529 | 0.38014 | -1.54 | 0.12591 |
| Edu | 0.14239 | 0.0571 | 2.494 | 0.01381 |
| FD | -2.2235 | 2.22851 | -0.998 | 0.31927 |
| **DAN** | 0.09702 | 0.03614 | 2.685 | **0.00774** |

**Supplementary Table S5.** Linear mixed effect models assessing relationships between the within-FCS of the default mode network (DMN) and MoCA

| IV: MoCA | Estimate *β* | SE | t | p |
| --- | --- | --- | --- | --- |
| Intercept | 26.11067 | 1.94461 | 13.427 | <0.0001 |
| Age | -0.02466 | 0.02859 | -0.862 | 0.3899 |
| Sex | -0.66481 | 0.38036 | -1.748 | 0.0827 |
| Edu | 0.14104 | 0.05709 | 2.47 | 0.0147 |
| FD | -1.07901 | 2.29225 | -0.471 | 0.6382 |
| **DMN** | 0.04627 | 0.02163 | 2.139 | **0.0334** |

**Supplementary Table S6.** Linear mixed effect models assessing relationships between the within-FCS of the visual network (Visual) and MoCA

| IV: MoCA | Estimate *β* | SE | t | p |
| --- | --- | --- | --- | --- |
| Intercept | 26.49836 | 1.96698 | 1.35E+01 | <0.0001 |
| Age | -0.0249 | 0.02904 | -0.857 | 0.3927 |
| Sex | -0.62198 | 0.38509 | -1.615 | 0.1085 |
| Edu | 0.14452 | 0.05789 | 2.496 | 0.0137 |
| FD | -1.96595 | 2.25093 | -0.873 | 0.3832 |
| Visual | 0.02675 | 0.01822 | 1.468 | 0.1433 |

**Supplementary Table S7.** Linear mixed effect models assessing relationships between the within-FCS of the ventral attention network (VAN) and MoCA

| IV: MoCA | Estimate *β* | SE | t | p |
| --- | --- | --- | --- | --- |
| Intercept | 26.94787 | 1.96496 | 13.714 | <0.0001 |
| Age | -0.02816 | 0.02894 | -0.973 | 0.332 |
| Sex | -0.63355 | 0.38554 | -1.643 | 0.1026 |
| Edu | 0.14362 | 0.0579 | 2.481 | 0.0143 |
| FD | -1.77082 | 2.3177 | -0.764 | 0.4455 |
| VAN | 0.02152 | 0.033 | 0.652 | 0.5148 |

**Supplementary Table S8.** Linear mixed effect models assessing relationships between the within-FCS of the limbic network (Limbic) and MoCA

| IV: MoCA | Estimate *β* | SE | t | p |
| --- | --- | --- | --- | --- |
| Intercept | 27.34847 | 1.89955 | 14.397 | <0.0001 |
| Age | -0.02905 | 0.02889 | -1.006 | 0.3163 |
| Sex | -0.62212 | 0.38498 | -1.616 | 0.1084 |
| Edu | 0.14288 | 0.05788 | 2.469 | 0.0148 |
| FD | -2.00487 | 2.30731 | -0.869 | 0.3856 |
| Limbic | -0.01611 | 0.06577 | -0.245 | 0.8067 |

**Supplementary Table S9.** Linear mixed effect models assessing relationships between the within-FCS of the frontoparietal control network (FPCN) and MoCA

| IV: MoCA | Estimate *β* | SE | t | p |
| --- | --- | --- | --- | --- |
| Intercept | 27.52925 | 1.92638 | 14.291 | <0.0001 |
| Age | -0.02962 | 0.02887 | -1.026 | 0.3065 |
| Sex | -0.62089 | 0.38448 | -1.615 | 0.1086 |
| Edu | 0.14299 | 0.05779 | 2.474 | 0.0146 |
| FD | -2.13641 | 2.25536 | -0.947 | 0.3443 |
| FPCN | -0.02234 | 0.03782 | -0.591 | 0.5553 |

$$\Delta Cognitive Score\approx1 + age +edu+ sex+ \Delta[Within\text{-}FCS]$$

**In TG dataset (*n* = 112; post-intervention – pre-intervention):**

**Supplementary Table S10.** Linear regression models assessing relationships between the Δwithin-FCS of the SMN and ΔMoCA

| IV: MoCA | Estimate *β* | SE | t | p |
| --- | --- | --- | --- | --- |
| Intercept | 1.31679 | 1.98724 | 0.663 | 0.509 |
| Age | -0.02347 | 0.03092 | -0.759 | 0.45 |
| Sex | 0.22356 | 0.39583 | 0.565 | 0.574 |
| Edu | 0.08431 | 0.0626 | 1.347 | 0.181 |
| SMN | -0.01595 | 0.02449 | -0.651 | 0.516 |

**Supplementary Table S11.** Linear regression models assessing relationships between the Δwithin-FCS of the DAN and ΔMoCA

| IV: MoCA | Estimate *β* | SE | t | p |
| --- | --- | --- | --- | --- |
| Intercept | 1.714172 | 1.845898 | 0.929 | 0.3555 |
| Age | -0.025432 | 0.028661 | -0.887 | 0.3772 |
| Sex | 0.001589 | 0.393472 | 0.004 | 0.9968 |
| Edu | 0.073179 | 0.061406 | 1.192 | 0.2364 |
| **DAN** | 0.100691 | 0.046023 | 2.188 | **0.0312** |

**Supplementary Table S12.** Linear regression models assessing relationships between the Δwithin-FCS of the DMN and ΔMoCA

| IV: MoCA | Estimate *β* | SE | t | p |
| --- | --- | --- | --- | --- |
| Intercept | 1.76892 | 1.88199 | 0.94 | 0.35 |
| Age | -0.03008 | 0.02914 | -1.032 | 0.305 |
| Sex | 0.12009 | 0.39671 | 0.303 | 0.763 |
| Edu | 0.08682 | 0.06233 | 1.393 | 0.167 |
| DMN | 0.02788 | 0.02581 | 1.08 | 0.283 |

**Supplementary Table S13.** Linear regression models assessing relationships between the Δwithin-FCS of the Visual and ΔMoCA

| IV: MoCA | Estimate *β* | SE | t | p |
| --- | --- | --- | --- | --- |
| Intercept | 1.52453 | 1.91888 | 0.794 | 0.429 |
| Age | -0.02754 | 0.02957 | -0.931 | 0.354 |
| Sex | 0.23452 | 0.3998 | 0.587 | 0.559 |
| Edu | 0.08732 | 0.06267 | 1.393 | 0.167 |
| Visual | -0.01513 | 0.02591 | -0.584 | 0.561 |

**Supplementary Table S14.** Linear regression models assessing relationships between the Δwithin-FCS of the VAN and ΔMoCA

| IV: MoCA | Estimate *β* | SE | t | p |
| --- | --- | --- | --- | --- |
| Intercept | 1.707768 | 1.896956 | 0.9 | 0.37 |
| Age | -0.029753 | 0.029435 | -1.011 | 0.315 |
| Sex | 0.19587 | 0.396581 | 0.494 | 0.623 |
| Edu | 0.085026 | 0.063024 | 1.349 | 0.181 |
| VAN | -0.003893 | 0.042005 | -0.093 | 0.926 |

**Supplementary Table S15.** Linear regression models assessing relationships between the Δwithin-FCS of the Limbic and ΔMoCA

| IV: MoCA | Estimate *β* | SE | t | p |
| --- | --- | --- | --- | --- |
| Intercept | 1.718167 | 1.893199 | 0.908 | 0.366 |
| Age | -0.030051 | 0.029322 | -1.025 | 0.308 |
| Sex | 0.190783 | 0.393663 | 0.485 | 0.629 |
| Edu | 0.086031 | 0.062833 | 1.369 | 0.174 |
| Limbic | -0.009565 | 0.089283 | -0.107 | 0.915 |

**Supplementary Table S16.** Linear regression models assessing relationships between the Δwithin-FCS of the FPCN and ΔMoCA

| IV: MoCA | Estimate *β* | SE | t | p |
| --- | --- | --- | --- | --- |
| Intercept | 1.737946 | 1.904372 | 0.913 | 0.364 |
| Age | -0.030216 | 0.029413 | -1.027 | 0.307 |
| Sex | 0.196523 | 0.397468 | 0.494 | 0.622 |
| Edu | 0.084988 | 0.063063 | 1.348 | 0.181 |
| FPCN | -0.004826 | 0.051865 | -0.093 | 0.926 |

$$Cognitive Score\approx1 + age +edu+ sex+ [Within\text{-}FCS]+headmotion$$

**Within TG2 dataset (*n* = 112):**

**Supplementary Table S17.** Linear regression models assessing relationships between the within-FCS of the SMN and MoCA after intervention trials

| IV: MoCA | Estimate *β* | SE | t | p |
| --- | --- | --- | --- | --- |
| Intercept | 28.33138 | 2.49875 | 11.338 | <0.0001 |
| Age | -0.05932 | 0.0406 | -1.461 | 0.1475 |
| Sex | -0.56261 | 0.53216 | -1.057 | 0.2932 |
| Edu | 0.19102 | 0.08222 | 2.323 | 0.0224 |
| FD | -0.11078 | 4.09089 | -0.027 | 0.9785 |
| SMN | 0.03663 | 0.03627 | 1.01 | 0.3152 |

**Supplementary Table S18.** Linear regression models assessing relationships between the within-FCS of the DAN and MoCA after intervention trials

| IV: MoCA | Estimate *β* | SE | t | p |
| --- | --- | --- | --- | --- |
| Intercept | 26.11687 | 2.5901 | 10.083 | <0.0001 |
| Age | -0.03932 | 0.03922 | -1.002 | 0.3188 |
| Sex | -0.58902 | 0.51602 | -1.141 | 0.2567 |
| Edu | 0.17626 | 0.07991 | 2.206 | 0.0299 |
| FD | -0.58581 | 3.96382 | -0.148 | 0.8828 |
| **DAN** | 0.17987 | 0.06872 | 2.617 | **0.0104** |

**Supplementary Table S19.** Linear regression models assessing relationships between the within-FCS of the DMN and MoCA after intervention trials

| IV: MoCA | Estimate *β* | SE | t | p |
| --- | --- | --- | --- | --- |
| Intercept | 25.65935 | 2.7137 | 9.455 | <0.0001 |
| Age | -0.03769 | 0.03959 | -0.952 | 0.3436 |
| Sex | -0.72456 | 0.52257 | -1.387 | 0.169 |
| Edu | 0.18524 | 0.08018 | 2.31 | 0.0231 |
| FD | 1.47744 | 4.05226 | 0.365 | 0.7163 |
| **DMN** | 0.10648 | 0.04418 | 2.41 | **0.018** |

**Supplementary Table S20.** Linear regression models assessing relationships between the within-FCS of the Visual and MoCA after intervention trials

| IV: MoCA | Estimate *β* | SE | t | p |
| --- | --- | --- | --- | --- |
| Intercept | 28.35844 | 2.6424 | 10.732 | <0.0001 |
| Age | -0.05248 | 0.04032 | -1.302 | 0.1963 |
| Sex | -0.57521 | 0.53487 | -1.075 | 0.285 |
| Edu | 0.19039 | 0.08264 | 2.304 | 0.0235 |
| FD | -0.22239 | 4.11792 | -0.054 | 0.957 |
| Visual | 0.01179 | 0.03979 | 0.296 | 0.7676 |

**Supplementary Table S21.** Linear regression models assessing relationships between the within-FCS of the VAN and MoCA after intervention trials

| IV: MoCA | Estimate *β* | SE | t | p |
| --- | --- | --- | --- | --- |
| Intercept | 25.87637 | 2.65176 | 9.758 | <0.0001 |
| Age | -0.05352 | 0.039 | -1.372 | 0.17331 |
| Sex | -0.57826 | 0.51756 | -1.117 | 0.26681 |
| Edu | 0.2215 | 0.08093 | 2.737 | 0.00746 |
| FD | 3.29031 | 4.2273 | 0.778 | 0.43838 |
| **VAN** | 0.16926 | 0.06766 | 2.502 | **0.01415** |

**Supplementary Table S22.** Linear regression models assessing relationships between the within-FCS of the Limbic and MoCA after intervention trials

| IV: MoCA | Estimate *β* | SE | t | p |
| --- | --- | --- | --- | --- |
| Intercept | 28.72908 | 2.49969 | 11.493 | <0.0001 |
| Age | -0.05176 | 0.04029 | -1.285 | 0.2021 |
| Sex | -0.58293 | 0.53437 | -1.091 | 0.2782 |
| Edu | 0.19368 | 0.08272 | 2.341 | 0.0214 |
| FD | 0.18253 | 4.18878 | 0.044 | 0.9653 |
| Limbic | -0.06171 | 0.10635 | -0.58 | 0.5632 |

**Supplementary Table S23.** Linear regression models assessing relationships between the within-FCS of the FPCN and MoCA after intervention trials

| IV: MoCA | Estimate *β* | SE | t | p |
| --- | --- | --- | --- | --- |
| Intercept | 29.31526 | 2.67146 | 10.973 | <0.0001 |
| Age | -0.05443 | 0.04026 | -1.352 | 0.18 |
| Sex | -0.56254 | 0.5337 | -1.054 | 0.295 |
| Edu | 0.19066 | 0.08244 | 2.313 | 0.023 |
| FD | -0.4361 | 4.10081 | -0.106 | 0.916 |
| FPCN | -0.06523 | 0.09039 | -0.722 | 0.472 |
